# Early life adversity is associated with mental health and life histories through short-term mindsets

**DOI:** 10.64898/2026.06.04.730040

**Authors:** Bence C. Farkas, Valentin Wyart, Pierre O. Jacquet

## Abstract

Early life adversity is associated with increased risk for psychopathology and shifts in life history (LH) strategies, but the psychological mechanisms underlying these associations remain unclear. Drawing on evolutionary-developmental theory, we examined whether short-term mindsets mediate associations between dimensions of childhood adversity and mental health and LH-related outcomes. In a UK-representative sample of 877 adults, we assessed threat, deprivation, and unpredictability, alongside internalizing and externalizing symptoms, borderline features, and a latent factor capturing reproductive versus somatic maintenance effort. Structural equation models showed that adversity predicted poorer mental health and faster LH strategies. Short-term mindsets, indexed by lower future orientation and higher affective impulsivity, mediated effects of adversity, particularly unpredictability. Further decomposition of unpredictability into short-timescale and long-timescale forms revealed dissociable effects, with short-timescale unpredictability primarily linked to psychopathology and long-timescale unpredictability to reproductive-oriented LH strategies.

## Introduction

Compared to nonhuman primates, the human life cycle is characterized by relatively earlier weaning leading to a longer period of dependence on resources procured by others, and slower growth (Bogin, 1990; Bogin & Smith, 1996; Walker et al., 2006). This unique developmental stage of childhood is thought to be crucial to the remarkable learning ability and capacity for complex culture our species exhibits (Bogin, 1997; Kaplan et al., 2000). This period is marked by a great degree of phenotypic plasticity, adjusting the phenotype to current and expected future environments (Frankenhuis & Nettle, 2020; West-Eberhard, 2003). This includes shaping behaviour, socioemotional (McLaughlin & Gabard-Durnam, 2022) and cognitive mechanisms (Kolb & Gibb, 2014), personality features (Beuchot et al., 2026) and neuroendocrinological processes (Ellison, 2017). Unfortunately, our susceptibility to environmental conditions during early life can also contribute to the development of undesirable mental and physical health outcomes, especially when these environmental conditions include adverse experiences (Wade et al., 2022). Early life adversity can be generally defined as negative childhood experiences, such as poverty, neglect, victimization or maltreatment, that require significant psychological and behavioural adaptation by a typical child (Frankenhuis & Amir, 2022; McLaughlin et al., 2019).

Meta-analytic evidence indicates that approximately 60% of individuals are exposed to at least one adverse childhood experience throughout their lifetime (Bellis et al., 2019; Madigan et al., 2023). A substantial literature shows that such early life adversity contributes to poorer outcomes in physical health (Felitti et al., 1998; Hughes et al., 2017; Rod et al., 2020), mental health (Green et al., 2010; Kessler et al., 2005; McLaughlin et al., 2010b, 2010a), and academic performance (Chang et al., 2019; Yeo et al., 2024). According to estimates by Kessler et al. (2010), the proportion of global lifetime DSM-IV/CIDI disorders statistically attributable to the presence of early life adversity is about 30%. Beyond the ethical imperative to reduce human suffering, childhood adversity also imposes considerable societal costs, with annual economic burdens estimated at approximately $581 billion in North America and $748 billion in Europe (Bellis et al., 2019). Accordingly, advancing our understanding of the ultimate and proximate mechanisms through which adversity shapes developmental outcomes – and identifying effective strategies to mitigate its impact – remains a critical priority for public health.

Earlier frameworks for conceptualizing and operationalizing early life adversity focused on either isolating specific risk factors and mechanisms, or on using a cumulative risk approach (Evans et al., 2013). More recently, dimensional models have been proposed as alternatives (McLaughlin et al., 2021). In these evolutionary-developmental frameworks, individual adversity types impact development through neither fully distinct nor fully overlapping mechanisms. Instead, adversity experiences are clustered into a set of core dimensions based on shared features and mechanisms. The harshness-unpredictability model draws on life history (LH) theory, and proposes that dimensions of environmental stress do not merely disrupt developmental trajectories, but adaptively calibrate them to increase biological fitness, potentially at the cost of individual health and well-being (Belsky et al., 2012; Ellis et al., 2009, 2024). The model identifies two fundamental dimensions of ecological adversity: harshness, referring to extrinsic mortality-morbidity, and unpredictability, referring to stochastic variability in extrinsic mortality-morbidity. Both dimensions are proposed to shape LH strategies, i.e., coordinated developmental patterns of physiological, morphological, and behavioural traits, and their psychological and neuroendocrine mediators (Del Giudice et al., 2015). Specifically, evidence suggests that early life adversity shifts developing individuals towards adopting ‘fast’ LH strategies once adults, that prioritize shorter-term reproductive goals, over longer-term goals related to somatic maintenance (Brumbach et al., 2009; Ellis et al., 2024; Gaydosh et al., 2018; Mell et al., 2018; Simpson et al., 2012; Szepsenwol et al., 2017). The adoption of such a strategy is facilitated by the development of an unpredictability schema, which refers to pervasive belief that the world is unpredictable, uncontrollable, and untrustworthy (Cabeza de Baca et al., 2016; Hill et al., 2008; Manczak et al., 2017; Ross & Hill, 2002). This belief can then pave the way for short-term mindsets: cognitive, affective, and behavioural tendencies that focus on present goals, while discounting future ones (Deitzer et al., 2025; Farkas et al., 2022; Pepper & Nettle, 2017; van Gelder et al., 2025). The threat-deprivation model also views early life adversity through an adaptive lens, but focuses on adaptation to current environmental contexts, instead of broader ecological ones (McLaughlin et al., 2019; Miller et al., 2021; Sheridan et al., 2017). It identifies two adversity dimensions, based on the way they influence neuroplasticity. Threat is defined as harm or threat of harm, and it seems to primarily impact experience-dependent neuroplasticity of socioemotional development in the fronto-amygdala circuit that leads to hypervigilance for environmental threat (McLaughlin et al., 2019; Pollak et al., 2000; Samaey et al., 2024; Saxbe et al., 2018; Weissman et al., 2020). Deprivation is defined as lack of expected environmental input, and it is generally found to impair experience-expectant learning mechanisms in frontocentral cortical regions, leading to cognitive and linguistic difficulties (Bouton et al., 2024; Hodel et al., 2015; McLaughlin et al., 2019; Miller et al., 2021; Sheridan et al., 2017). The most recent dimensional model (Ellis et al., 2022) integrates these two earlier approaches, and considers threat (harshness stemming from danger), deprivation (harshness stemming from insufficient environmental input), and unpredictability as the three fundamental dimensions of early life adversity, with a growing literature empirically supporting the validity of this model (Farkas & Jacquet, 2024; Lee et al., 2024; Shaul et al., 2024; Usacheva et al., 2022).

In this study, we investigate the associations between dimensions of childhood adversity, mental health and LH traits, and test whether they are mediated by short-term mindsets. We also examine the dimension of unpredictability in more detail, given that it is often found to be even more strongly linked to developmental outcomes than harshness, and at the same time, its conceptualization and measurement remain challenging (Martinez et al., 2022; Young et al., 2020, 2022). There is ample empirical evidence supporting developmental cascade introduced above in which early life unpredictability leads individuals shift to a LH strategy characterized by increased reproductive over maintenance effort by adopting an unpredictability schema and a short-term mindset, leading to risk taking, impulsivity, and externalizing behavioural problems (Cabeza de Baca et al., 2016; Deitzer et al., 2025; Doom et al., 2016; Farkas et al., 2024; Farkas & Jacquet, 2024; Griskevicius et al., 2011, 2013; Martinez et al., 2022; Szepsenwol et al., 2017). However, the literature has not yet reached consensus on how to define and operationalize unpredictability as a construct. Researchers tend to either adopt an ‘ancestral cue’ approach, in which children are assumed to be sensitive to a limited set of specific experiences that served as cues to expected environmental variability in our evolutionary history, or a ‘statistical learning’ approach, which sees children as continuously estimating variability in their environment by more domain-general learning mechanisms (Z. Li et al., 2023; Young et al., 2020). As a result, quite diverse experiences, such as residential changes (e.g., Belsky et al., 2012), subjective retrospective judgments of perceived unpredictability (e.g., Jonason et al., 2016), inconsistent parenting (e.g., Wuth et al., 2021), family chaos (e.g., Evans et al., 2005), or income variability (Zachrisson & Dearing, 2015) have all been used as proxies of unpredictability. There are some findings suggesting that the timescale and type of uncertainty might be crucial in determining its developmental impact (Chang et al., 2021; Cohodes et al., 2023; DeJoseph et al., 2025; Farkas et al., 2024; Hartman et al., 2018; Walasek et al., 2024). Here, we tentatively adopt an organizing principle which distinguishes between short-timescale and long-timescale unpredictability. This is partially inspired by the human reinforcement learning (RL) literature, which distinguishes between ‘stochastic’ and ‘volatile’ forms of uncertainty, with important consequences for adaptive behaviour (Piray & Daw, 2021; Soltani & Izquierdo, 2019). Stochasticity, or ‘expected uncertainty’, involves random noise around a stable mean (e.g., daily variation in commute time when using public transport), while volatility, or ‘unexpected uncertainty’, signals a true change in outcome contingencies (e.g., a sudden subway closure). There is extensive empirical evidence that humans can arbitrate between these different forms of uncertainty in their environment, and adopt learning and decision-making styles in an adaptive manner, depending on the degree of inferred volatility and stochasticity (Behrens et al., 2007; J. K. Lee et al., 2023; Piray & Daw, 2024; Soltani & Izquierdo, 2019). Early life experiences might also be subject to uncertainty classifiable along similar dimensions. For example, day-to-day variability in the time parents come back home is more akin to stochastic unpredictability, whereas an abrupt residential or school change seems closer to volatile unpredictability (Farkas et al., 2024). Building on this, we also explore whether distinguishing relatively more stochastic, short-timescale from more volatile, long-timescale early life unpredictability experiences has utility for explaining variation in mental health and LH outcomes.

We know of no previous study that thoroughly investigated the mediating role of short-term mindsets in the evolutionary-developmental framework using both a state-of-the-art dimensional assessment of childhood adversity and an approximation of individual LH strategies using biodemographic markers. To fill this gap, we recruited a sample of 877 adults, representative of the UK population through an online data collection platform. We used a structural equation modelling framework to investigate associations between retrospectively reported early life threat, deprivation, and unpredictability on general mental health functioning (including internalizing symptoms, externalizing problems, and borderline personality traits), somatic maintenance effort and reproductive effort, and whether they are mediated by short-term mindsets, while adjusting for confounds (age, ethnicity, sex, current socio-economic status). In addition, we explored whether the effects of unpredictability can be better accounted for by separate short-timescale (ST) and long-timescale (LT) constructs.

## Methods

### Ethics and open science

The research was carried out following the principles and guidelines for experiments including human participants provided in the Declaration of Helsinki (World Medical Association, 2025). The study was approved by the local ethical committee (N°CERES 201659). All anonymised data, analysis code, and materials for this study are publicly available on the Open Science Framework (OSF) repository at the following link : https://osf.io/yzp2x. The study was not pre-registered.

### Participants

We aimed to recruit a representative sample on Prolific (prolific.co; Palan & Schitter, 2018), targeting the United Kingdom (UK). UK representative samples on Prolific are stratified based on age into five brackets: 18-24, 25-34, 35-44, 45-54 and 55+; based on sex into male and female; and based on ethnicity into the five categories recommended by the UK Office of National Statistics: White, Mixed, Asian, Black and Other. These variables are then matched to the UK census. For more information on Prolific representative samples see https://researcher-help.prolific.com/en/articles/445161-what-are-representative-samples-on-prolific.

Minimum sample size was determined through an *a priori* model-based power analysis of our focal hypothesis using the *semPower* R package (Moshagen & Bader, 2023). This method works by translating an effect to be tested in a user-specified structural equation model into the expected effect on the population and model-implied covariance matrices, and eventually the population minimum of the fit function *F_0_*. We carried out power analysis based on a simplified version of our analytic model, with population values based on relevant earlier studies (Cabeza de Baca et al., 2016; Deitzer et al., 2025; Farkas & Jacquet, 2024; Martinez et al., 2022). Specifically, we tested whether the indirect effect of early life unpredictability (observed variable) on the reproduction/maintenance latent (7 indicators with assumed loadings of λ = .4) through short-term mindset (2 indicators with assumed loadings of λ = .4) differs from zero in a simple latent mediation model. The population value was set to β = 0.20 for the early life unpredictability → short-term mindset (XM) path; to β = 0.20 for the short-term mindset → reproduction/maintenance (MY) path; and to β = 0.25 for the early life unpredictability → reproduction/maintenance (XY) path. This corresponds with a modest standardized direct effect of β = 0.25, and a small standardized indirect effect of β = 0.04. This analysis suggested a minimum sample size of 720 is required to detect the indirect effect with 80% power, at α = .05.

Taking into account attrition, we recruited a representative sample of 1001 UK participants. Of these, 988 had completed the study fully. One participant was excluded for failing an attention check question, 94 participants were excluded for reporting implausible values on variables (e.g., reporting no sexual intercourse yet, but reporting nonzero number of sexual partners), and 20 participants were excluded for reporting the presence of a serious neurological disorder (e.g., epilepsy, brain injury, stroke, multiple sclerosis). Note that we did not exclude participants for reporting other mental health or neurodevelopmental conditions. This led to a final sample size of 877 (participants could meet more than one exclusion criterion). This is well above the minimum sample size suggested by our power analysis (N ≥ 720), and above rules-of-thumb thresholds for structural equation models (N ≥ 450; Wolf et al., 2013), and reliability analyses (N ≥ 200; Streiner et al., 2024).

### Measures

#### Early life adversity

We aimed to capture participants exposure to multiple dimensions of early life adversity, as outlined in the prevailing dimensional framework. Accordingly, we measured early life threat (experiences that confer the risk of physical and psychological harm), deprivation (experiences that reflect insufficient environmental input) and unpredictability (experiences that signal spatiotemporal variability in deprivation and threat), all using recent, psychometrically validated scales. To equalize the relevant age range for our different measures, we modified the instructions to always inquire about experiences that occurred before age 12.

##### Early life deprivation and threat

We measured early life deprivation and threat with the Deprivation and Threat – Adult Self-report (DT-AS) scale (Tsai et al., 2025). The DT-AS is a comprehensive, self-report early adversity measure, that covers the dimensions of threat, and deprivation. It meets COSMIN criteria for high-quality health outcome measures (Mokkink et al., 2010), and demonstrated excellent psychometric properties in an initial validation study (Tsai et al., 2025). Items inquire about the frequency with which participants experienced a given event during childhood, and are rated on a Likert scale with options: 1-Never (coded as 0), 2 - Almost never (coded as 1), 3 - Sometimes (coded as 2), 4 - A lot (coded as 3). Some items are reverse coded. The Deprivation subscale consists of 30 items, whereas the Threat subscale consists of 33 items. Example items include ‘How often did an adult show an interest in your thoughts and feelings at home?’ (Deprivation) and ‘How many times has an adult pushed, grabbed, slapped, or thrown something at you at home?’ (Threat). Participants final scores are the unweighted averages of the ratings in the original paper, but we used the sum score instead for easier interpretation. We made two important modifications to the original measure. First, the original measure uses a timeframe of ‘prior to age 18’ for all items. In order to focus on the pre-adolescent developmental period that is expected to be the most impactful in organizing developmental trajectories (Gee & Cohodes, 2021; Simpson et al., 2012), we homogenised the scale, such that in our implementation all questions were phrased so as to refer to the period ‘prior to age 12’. Second, in order to incorporate individuals’ subjective perception of adverse experiences (Danese & Widom, 2020; Lacey & Minnis, 2020), after each item, we added a further question that inquired about the subjective impact of the given experience (“How did you feel about this at the time?”), with response options: 1 - Very negatively, 2 - Negatively, 3 - No strong feelings / I don’t know, 4 - Positively, 5 - Very positively. In the main analysis, only the main deprivation and threat scale scores are used as indices of early life deprivation and threat exposure, with higher scores indicating greater exposure. The deprivation scale has a potential range = [0, 90], and observed scores had a median of 25, with a median absolute deviation of 14.83, and range = [2,78] (Supplementary Figure S1). It demonstrated excellent internal consistency in the sample, Cronbach α = .92 [.91, .93]; average inter-item r = .29. The threat scale has a potential range = [0, 99], and observed scores had a median of 14.5, with a median absolute deviation of 11.12, and range = [0,87] (Supplementary Figure S1). It also demonstrated excellent internal consistency in the sample, Cronbach α = .93 [.92, .94]; average inter-item r = .28.

##### Early life unpredictability

We measured early life unpredictability with the Questionnaire of Unpredictability in Childhood (QUIC) - a self-report measure designed to assess lack of predictability in multiple aspects of the childhood environment (Glynn et al., 2019). The scale has demonstrated strong internal consistency, excellent test–retest reliability and has been shown to predict other unpredictability indicators and internalizing symptoms, suggesting good construct validity. Items are binary, with ‘yes’ (coded as 1) or ‘no’ (coded as 0) response options, with some items reverse coded. Example items include ‘At least one of my parents was unpredictable’ and ‘At least one of my parents regularly checked that I did my homework’. We made the same two changes to the original questionnaire as for the DT-AS. First, the original questionnaire uses a timeframe of ‘prior to age 12’ for some items and ‘prior to age 18’ for others. We modified this, such that in our implementation all questions were phrased so as to refer to the period ‘prior to age 12’. Second, we added the same question about the subjective impact of the given experience. Contrary to the DT-AS, the QUIC does not refer to frequencies, which makes responding to the impact questions after non-endorsed items difficult. As a result, for the QUIC, we only asked the impact questions after endorsed items (for reverse coded items, endorsement is the ‘no’ answer). In the main analysis, following earlier studies, we used the total sum score as an index of early life unpredictability exposure, with higher scores indicating greater exposure. This unpredictability scale has a potential range = [0, 38], and observed scores had a median of 6, with a median absolute deviation of 7.41, and range = [0,33] (Supplementary Figure S1). This score demonstrated excellent internal consistency in the sample, Cronbach α = .91 [.90, .92]; average inter-item r = .20.

#### Short-term mindsets

We aimed to assess both the cognitive aspect of short-term mindsets, reflecting present oriented thinking, and the affective/behavioural component, reflecting short-sighted emotional responses, and their tendency to elicit impulsive actions. To this end, we used future orientation as our proxy for the former, and affective impulsivity as our proxy for the latter component.

##### Future orientation

We measured future orientation with the Future Orientation Scale (FOS), a 15-item self-report measure of future orientation (Steinberg et al., 2009). This scale uses a format developed by Harter (1982) to reduce socially desirable responding. Participants are presented with 10 pairs of contrasting statements separated by the conjunction “BUT” and asked to select the option that best describes them. They then indicate whether the chosen statement is “really true” or “sort of true.” Responses are coded on a 4-point Likert scale reflecting the strength and direction of endorsement across the paired statements and averaged. The scale can also be subdivided into three, 5-item subscales for time perspective, anticipation of future consequences, and planning ahead. Example items include ‘A. Some people like to plan things out one step at a time. BUT B. Other people like to jump right into things without planning them out beforehand.’ and ‘A. Some people like to think about all of the possible good and bad things that can happen before making a decision. BUT B. Other people don’t think it’s necessary to think about every little possibility before making a decision.’. In the main analysis, we only used the overall future orientation score as an index of future orientation, with higher scores reflecting greater future orientation / less present orientation. This scale has a potential range = [1, 4], and observed scores had a median of 3, with a median absolute deviation of 0.49, and range = [1,4] (Supplementary Figure S1). This score demonstrated good internal consistency in the sample, Cronbach α = .85 [.83, .86]; average inter-item r = .27.

##### Affective impulsivity

We measured affective impulsivity with the Feelings Trigger Action (FTA) subscale of a three-factor impulsivity index developed through factor analysis of existing scales by Carver et al. (2011). While reliability for the factor analytically derived subscales has never been published, the original instruments demonstrated excellent reliability, and the resulting subscales possess excellent construct validity (Carver et al., 2011; Hooper & Carver, 2016; Johnson et al., 2013). The FTA subscale has demonstrated transdiagnostic associations with internalizing, externalizing, and Borderline problems (Johnson et al., 2013). This 26-item scale is made up of items from an (negative) urgency instrument (Whiteside & Lynam, 2001), a positive urgency instrument (Cyders et al., 2007), and custom items tapping into reflexive reactions to feelings (Carver et al., 2011). Items are rated on a 5-point Likert scale, with the total score being the unweighted average of all items. For the main analysis, we used the total score, with higher scores reflecting a greater degree of affective impulsivity. This scale has a potential range = [1, 5], and observed scores had a median of 2.57, with a median absolute deviation of 1.03, and range = [1, 4.89] (Supplementary Figure S1). This score demonstrated excellent internal consistency in the sample, Cronbach α = .97 [.96, .97]; average inter-item r = .52.

#### Reproductive and maintenance effort

The trade-off between reproductive and somatic maintenance effort was modelled as a latent factor aiming to capture the common variance of seven indicators. Reproductive effort was indexed by four indicators focusing on the timing of reproductive effort, on fertility, and on preference for promiscuous sex, while maintenance effort was indexed by three indicators informing about physiological and behavioural factors that contribute to the participants’ general health state (Baptista et al., 2023; Farkas et al., 2022; Lettinga et al., 2021; Mell et al., 2018; Sýkorová & Flegr, 2021). Descriptive statistics and histograms for all untransformed indicators are presented in Supplementary Figure S1.

##### Sexual debut

We asked participants to report their age in years at their first consented sexual intercourse (Mell et al., 2018; Moffitt et al., 1992; Simpson et al., 2012). To not lose information from participants who have not had sexual intercourse yet, we binned this variable to four relatively equally sized bins (1 - earliest to 4 - latest), and assigned such individuals to bin 5. This variable was thus an ordered variable reflecting participants’ relative sexual debut in this sample: 1 (< 16y, N = 279, 31.8%), 2 (16y - 18y, N = 264, 30.1%), 3 (18y - 19y, N = 77, 8.8%), 4 (> 19y, N = 190, 21.7%), 5 (not yet, N = 67, 7.6%).

##### Number of sexual partners

We asked participants to report their total lifetime sexual partners (Lettinga et al., 2021; Mell et al., 2018). This variable was z scored before being entered into the model to mitigate potential issues arising from its large range.

##### Age at 1^st^ reproduction

We asked participants to report their age in years when their first child was born (Ellis et al., 2024; Mell et al., 2018; Tropf et al., 2015). We recoded this variable in the same manner as asexual debut for the same reason. As a result, age at 1^st^ reproduction was also an ordered variable reflecting participants’ relative age at first childbirth in this sample: 1 (< 24y, N = 138, 15.7%), 2 (24y - 28y, N = 114, 13.0%), 3 (28y - 32y, N = 106, 12.1%), 4 (> 32y, N = 116, 13.2%), 5 (not yet, N = 403, 46.0%).

##### Number of children

We asked participants to report their total number of biological children (Lettinga et al., 2021; Mell et al., 2018). Note that participants could have an age at 1^st^ reproduction, but a response of 0 for number of children, if they lost their child. This variable was recoded by collapsing individuals with 3 or more children into a single category. Thus, it was treated as ordered with the following categories: 0 (N = 448, 51.1%), 1 (N = 130, 14.8%), 2 (N = 198, 22.6%), 3 (N = 101, 11.5%).

##### Health status

Participants were asked to make a subjective report of their general state of health (from 1-‘Very poor’ to 6-‘Excellent’). Self-rated health is known to predict morbidity (Barger & Muldoon, 2006; Goldberg et al., 2001) and mortality (Benyamini & Idler, 1999; Idler & Benyamini, 1997), over and above traditional physiological risk factors (Haring et al., 2011; Jylhä et al., 2006).

##### Health efforts

Participants were also asked to report how much effort they make to look after their health and ensure their safety these days (from 1-‘None at all’ to 5-‘A great deal’). Questions about motivation towards looking after health is associated with a range of health behaviours (Becker et al., 1972; Mirotznik et al., 1998).

##### Smoking

Participants were asked ‘In the last 30 days, on average, how many cigarettes have you smoked per week?’. Alcohol and tobacco consumption are common detrimental health behaviours, that is associated with both childhood and adult stress, and thought to reflect present-oriented decision-making (Legleye et al., 2011; Mell et al., 2018; Melotti et al., 2011; Pepper & Nettle, 2017). This variable was also z scored before being entered into the model to mitigate potential issues arising from its large range.

#### Mental health

We opted for a transdiagnostic and dimensional assessment of general mental health, given its alignment with evolutionary-developmental psychopathology’s theoretical assumptions and empirical findings, and the non-clinical nature of the sample. There is extensive evidence that symptoms of psychopathology can be usefully organized into internalizing (mood and anxiety disorders) and externalizing (conduct disorder, ADHD, substance use and most personality disorders) spectra (Kessler et al., 2011; Krueger et al., 1998; Martel, 2013; Martel et al., 2017; Ringwald et al., 2021). However, some disorders defy a straightforward assignment into either spectrum. For example, Borderline Personality Disorder (BPD) appears to be composed of both externalizing and internalizing features (Eaton et al., 2011; Hudson et al., 2014; Wang et al., 2024). As BPD has featured extensively in life history-inspired psychopathology frameworks as a prototypical ‘fast’, reproductive effort maximizing condition (Baptista et al., 2023; Brüne, 2016; Del Giudice, 2018), we sought to assess Borderline features in more detail. To this end, we make use of a standard public mental health questionnaire for the general population covering the internalizing and externalizing spectra, and a more focused questionnaire developed to screen for Borderline features.

##### Internalizing and externalizing problems

We used the adult (17+) version of the Strengths and Difficulties Questionnaire (SDQ) to assess internalizing and externalizing problems (Brann et al., 2018). The adult SDQ is a 25-item self-report measure, suitable for measuring both overall psychological distress and more specific domains. Example items include ‘I worry a lot’ and ‘I get very angry and often lose my temper’. Some items are reverse coded. The SDQ was originally developed for children and adolescents, but an adult version, which shares the same structure and shows similarly good psychometric properties has been recently developed and validated (Armitage et al., 2023; Brann et al., 2018; Findon et al., 2016; Liang et al., 2025; Somma et al., 2020; Stringer et al., 2020). We preferred this questionnaire over the similar but more extensive Adult Self Report (Achenbach & McConaughy, 2003) due to its time and cost effectiveness, and better alignment with Open Science practices. Participants rated the items on a 3-point Likert scale (0 = Not true, 1 = Somewhat true, 2 = Certainly true). For the main analysis, we used the sum scores for the 10-item internalizing and externalizing subscales, with greater scores indicating greater internalizing and externalizing problems. The internalizing scale has a potential range = [0, 20], and observed scores had a median of 6, with a median absolute deviation of 4.45, and range = [0,18] (Supplementary Figure S1). This score demonstrated good internal consistency in the sample, Cronbach α = .80 [.78, .82]; average inter-item r = .28. The externalizing scale also has a potential range = [0, 20], and its observed scores had a median of 4, with a median absolute deviation of 2.97, and range = [0,17] (Supplementary Figure S1). The externalizing score demonstrated acceptable internal consistency in the sample, Cronbach α = .73 [.70, .76]; average inter-item r = .21.

##### Borderline features

We used the McLean Screening Instrument for Borderline Personality Disorder (MSI-BPD) to assess the presence of BPD features. The MSI-BPD is a 10-item self-report scale based on the borderline module of the Diagnostic Interview for DSM-IV Personality Disorders (Zanarini et al., 1996). The measure was originally developed as a screening tool, with the cutoff value of 7 leading to good sensitivity and specificity in detecting the presence of BPD across a wide range of samples (Keng et al., 2025; Mirkovic et al., 2020; Zanarini et al., 2003; Zimmerman & Balling, 2021). In addition, studies also confirmed the suitability of the measure for interindividual differences research by demonstrating excellent reliability and construct validity (Keng et al., 2025; Mirkovic et al., 2020). All items are binary (0 = No, 1 = Yes), with endorsement indicating the presence of a BPD feature. Example items include ‘Have you often been distrustful of other people?’ and ‘Have you chronically felt empty?’. For the main analysis, we used the sum score, with greater scores indicating more BPD features. This scale thus has a potential range = [0, 10], and observed scores had a median of 3, with a median absolute deviation of 4.45, and range = [0,10] (Supplementary Figure S1). This score demonstrated good internal consistency in the sample, Cronbach α = .86 [.84, .87]; average inter-item r = .37.

#### Adjustment variables

##### Current subjective socio-economic status

To assess subjective current socio-economic status (SES), we used the MacArthur Scale of Subjective Social Status (Adler et al., 2000), a single-item measure of an individual’s perceiver rank relative to others in their group, with 10 response options ranging from 1 (worst off) to 10 (best off). Higher values thus indicate lower perceived subjective social status. The median response was 5, with a median absolute deviation of 1.48, and range = [1,10] (Supplementary Figure S1).

##### Sex

We used the Sex variable from Prolific’s demographic database. It was coded as 0 = Female, 1 = Male for the structural equation models.

##### Ethnicity

We used the Ethnicity variable from Prolific’s demographic database, which contains the following categories: White, Mixed, Asian, Black and Other. We recoded this variable to 0 = White, 1 = Non-white for use as an exogenous adjustment variable in the models.

##### Age

We used the Age (in years) variable from Prolific’s demographic database.

### Analytical plan

Our analytic plan consists of four steps. First, we construct a model to investigate direct and indirect effects of early life deprivation, threat and general unpredictability on outcomes. Second, we split the general unpredictability score into more specific short (more stochastic)-and long (more volatile) timescale components and validate these novel subscales. Third, we construct a model with these more specific unpredictability scores to investigate their potentially distinct effects. Fourth, we fit two sensitivity models: the first one replaces our reproductive/maintenance trade-off latent construct by sexual debut only, and the second one replaces the early adversity scores with versions that weigh individual experiences by their subjective impact. These steps are described in more detail next.

#### ST/LT scale validation

One objective of the current study was to investigate early life unpredictability with more precision. Inspired by earlier work (DeJoseph et al., 2025; Farkas et al., 2024; Walasek et al., 2024) and the RL literature (Behrens et al., 2007; J. K. Lee et al., 2023; Piray & Daw, 2021; Soltani & Izquierdo, 2019), we explored whether this umbrella construct can be usefully subdivided, based on the type of uncertainty that a given experience is likely to signal. For the reasons outlined in the introduction, we focused on using the existing QUIC instrument to approximate the constructs of early life short-timescale (ST) and long-timescale (LT) unpredictability.

Our goal was to classify the items as reflecting either early life ST or LT unpredictability. To this aim, we used two complementary classification procedures: a theory-based, top-down classification procedure elaborated by us, complemented by a theory-free, bottom-up classification procedure performed by large language models. In our theory-based approach, for an item to be classify as ST or LT, its formulation must explicitly mention the occurrence of a short-timescale or long-timescale change in the subjects’ early family, social and physical environment. Three items from the Physical environment subscale of the QUIC questionnaire were discarded because their formulation referred to only spatial cluttering, rather than temporal variability (“I lived in a clean house” ; “I lived in a cluttered house (e.g. piles of stuff everywhere)” ; “In my house things I needed were often misplaced so that I could not find them”). The three items of the Safety and security subscale were also discarded as they seemed to measure physical deprivation and threat instead of unpredictability (“There was a period of time when I often worried that I was not going to have enough food to eat.” ; “There was a period of time when I often worried that my family would not have enough money to pay for necessities like clothing or bills.” ; “There was a period of time when I did not feel safe in my home.”). From the remaining items, eleven items were classified as reflecting LT unpredictability (e.g., “There was a long period of time when I did not see one of my parents – e.g., military deployment, jail time, custody arrangements”), and twenty-one were classified as reflecting ST unpredictability (e.g., “One of my parents could go from calm to furious in an instant”). The complete list of items and their classification can be found in the Supplementary Table S1. For the LLM item classification analysis, we used ChatGPT 5.3 (OpenAI, 2026) and Claude Sonnet 4.6 (Anthropic, 2026) using a web interface. There were three prompt conditions: i) no specific information; ii) a section from the introduction of a recent RL study on stochasticity and volatility (Piray & Daw, 2024); iii) a section from the introduction of our study. Full prompts can be found in Supplementary Text S1. Each model / prompt combination was performed 5 different times, with randomly permuted item order. Classification overlap was quantified with simple accuracy, and the Cohen’s κ.

#### Structural equation models

We tested the direct and indirect effects of early life adversity dimensions on our outcomes using latent mediation structural equation models (SEMs). Early life adversity variables were continuous manifest predictors, and mental health variables were continuous manifest outcomes. Short-term mindsets were modelled as a latent variable with two indicators, the affective impulsivity (FTA) and the future orientation (FOS) scales, and it served as our mediating variable. The trade-off between reproductive and maintenance effort was also considered as an outcome, and was modelled as a latent variable with four reproduction indicators (Sexual debut, Age at 1^st^ reproduction, Number of sexual partners, Number of children) and three maintenance indicators (Health status, Health efforts, Alcohol consumption, Smoking). We allowed for residual covariances between some indicators, that we expected to be correlated for reasons outside the hypothesized reproduction/maintenance trade-off. We allowed correlations between the health efforts and health status indicators, the age at 1^st^ reproduction and number of children indicators, and the sexual debut and number of sexual partners indicators. In addition, we added a residual covariance between the sexual debut and age at 1^st^ reproduction and number of children indicators based on modification indices. We specified direct effects between all early life adversity variables and all outcomes, as well as the corresponding indirect effects through short-term mindsets. We also specified additional residual correlations between the health status and health efforts indicators and the mental health outcomes, as we might expect them to covary negatively for reasons independent of the structural paths. Finally, all manifest variables were adjusted for age, ethnicity, sex, and current subjective SES.

Models were fit in *R* 4.5.2. (R Core Team, 2025) with the *lavaan* (Rosseel, 2012) and *semTools* (Jorgensen et al., 2020) packages, with the *tidyverse* package (Wickham et al., 2019) used for preprocessing and plotting. As the variables health status, health efforts, sexual debut, number of children, and age at 1^st^ reproduction were treated as ordered, the WLSMV estimator was used with delta parametrization, due to its robustness against violations of normality and ability to handle categorical manifest variables (C.-H. Li, 2016). Latent variables were scaled by fixing their variance to 1. There were no missing values and no outliers were removed, except for clearly implausible values (see the Participants section above). Parameters throughout the manuscript are presented in standardized form. Inference regarding direct and indirect effects was done via Monte Carlo confidence intervals (Preacher & Selig, 2012). All p values reported are two-tailed, with alpha set to .05. Goodness of fit was evaluated using the (scaled) Chi-squared test, comparative fit index (CFI), standardized root mean square residual (SRMR), root mean squared error of approximation (RMSEA) values. Given the Chi-squared tests sensitivity to sample size, relatively more weight was given to the other indices (Fabrigar et al., 1999; MacCallum, 1990). A commonly used heuristic from simulation studies suggests values of CFI ≥ .95, SRMR ≤ .08, RMSEA ≤ .06 is indicative of good fit (Hu & Bentler, 1999), with earlier proposed thresholds for CFI and RMSEA being somewhat more liberal at CFI ≥ .90 (Bentler & Bonett, 1980), and RMSEA ≤ .08 (Browne & Cudeck, 1993) suggested to indicate acceptable fit. However, to complement the weaknesses of fixed thresholds, we also evaluate the full residual matrix, which gives a more detailed picture of fit (Kline, 2023). Additionally, we use the *cvsem* package (Wysocki et al., 2022) to obtain 10-fold cross-validation indices for comparing the unpredictability and the short/long SEMs. This method quantifies cross-validation performance as the Kullback-Leibler (KL) divergence between the model implied covariance matrix estimated from the training data and the sample covariance matrix estimated from the test data, with lower values indicative of better performance.

We primarily rely on the differential association of the ST/LT subscales with the outcome variables, as evidence for their validity. We assessed this in a number of different ways. First, we qualitatively compared the fit indices and cross-validation performance of the unpredictability model and the ST/LT model. As the models are non-nested, and not fit with ML estimation, their direct statistical comparison is not possible. Second, we conducted a Wald test of linear equality constraints within the fitted ST/LT SEM (Wald, 1939). Specifically, all regression coefficients for the two predictors were constrained to be equal and then jointly tested against the null hypothesis that these equality constraints hold in the population. Third, we compared the ST/LT model to its constrained version, in which corresponding regression coefficients were set equal, with a chi-square difference test, evaluating whether imposing the equality constraints leads to a significant deterioration in overall model fit (Satorra, 2000).

## Results

We fit two latent mediation SEMs to a representative sample of 877 UK participants, in order to test four main hypotheses.

In a first general unpredictability SEM, we test the first three hypotheses:

H1) The covariance of reproductive and maintenance indicators will be captured by a latent variable reflecting a trade-off between reproductive and maintenance effort.

H2) Early life adversity, especially the dimension of threat, will have positive direct effects on all three mental health outcomes, as well as increase reproductive over maintenance effort.

H3) Early life adversity, especially the dimension of unpredictability, will have positive indirect effects on all three mental health outcomes, as well as increase reproductive over maintenance effort through increasing short-term mindsets.

In a second ST/LT SEM, we test the final hypothesis:

H4) A model separating the early life ST and LT unpredictability will show qualitatively better fit than a general unpredictability model, and will result in evidence for specific and dissociable effects for the two timescales of early life unpredictability.

Demographic information about the sample is presented in Table 1, distributions of pre-transformed variables are presented in Supplementary Figure S1, bivariate correlations between variables are presented in Figure 1. Correlations generally aligned with the hypotheses outlined above. Reproductive and maintenance effort indicators had small to medium sized negative correlations in directions consistent with a trade-off. They also covaried with early life adversity dimensions and mental health in the expected manner. Affective impulsivity and future orientation covaried negatively with each other, and both were related to all early life adversity dimensions, and most mental health scores.

**Figure 1.**
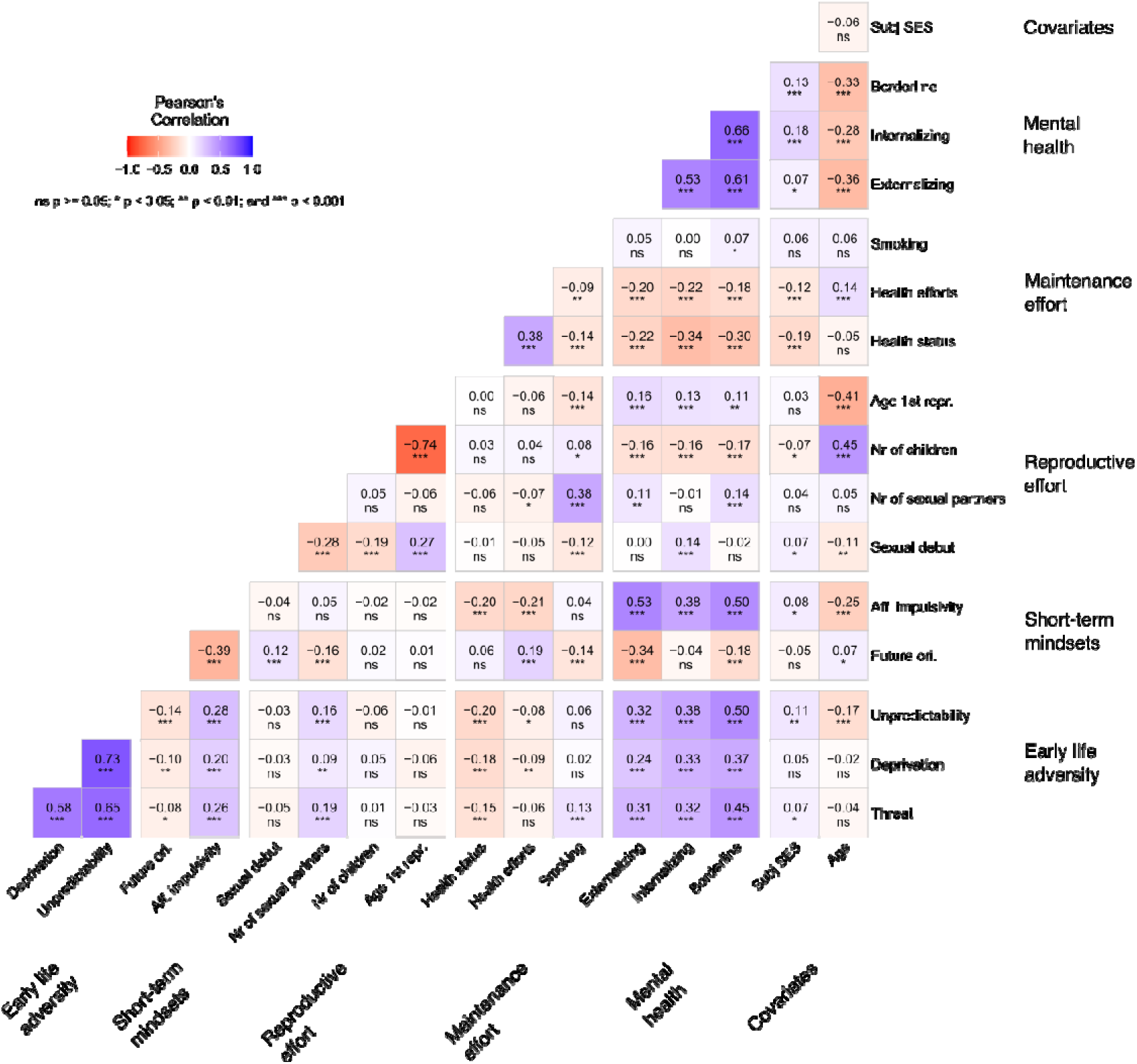
Bivariate correlations between study variables. Numbers are Pearson’s linear correlation coefficients. P values are uncorrected for multiple comparisons.

**Table 1.** Sample descriptive statistics.

|  | **M (SD)** | **N (%)** |
| --- | --- | --- |
| Age (years) | 46.13 (15.69) |  |
| Sex |  |  |
| Male |  | 426 (48.58%) |
| Female |  | 451 (51.42%) |
| Ethnicity |  |  |
| White |  | 749 (85.40%) |
| Mixed |  | 16 (1.82%) |
| Asian |  | 72 (8.21%) |
| Black |  | 25 (2.85%) |
| Other |  | 15 (1.71%) |
| Education |  |  |
| Some high school or less |  | 15 (1.71%) |
| Completed high school |  | 163 (18.59%) |
| Trade/technical school |  | 56 (6.39%) |
| Some college |  | 96 (10.95%) |
| Completed college |  | 339 (38.65%) |
| Post graduate |  | 208 (23.72%) |

### General unpredictability SEM

#### Model fit

The general unpredictability model presented with the following fit indices: χ^2^(57) = 285.94, p < .001; CFI = .942; RMSEA = .068 [.060, .076]; SRMR = .058. The model failed the Chi squared test, indicating poor fit, however the test is known to be sensitive to sample size. The CFI exceeds the liberal threshold of .90 indicating acceptable fit, but falls somewhat short of the conservative one of .95. The point estimate of RMSEA being below .08 also indicates acceptable fit, and the hypothesis of it falling below the .06 threshold for close fit cannot be rejected. The SRMR also indicates good fit. Standardized residuals (Supplementary Table S2) indicate generally tolerable levels of misfit (r < .10), with the exception of some unmodelled covariation between the health status and health efforts and the short-term mindset indicators and between internalizing symptoms and sexual debut (r ≈ .15 - .18). Overall, we deemed this model acceptable and retained it. A simplified diagram of the model is presented in Figure 2A, and the full table of fitted parameters is available in Supplementary Table S4.

**Figure 2.**
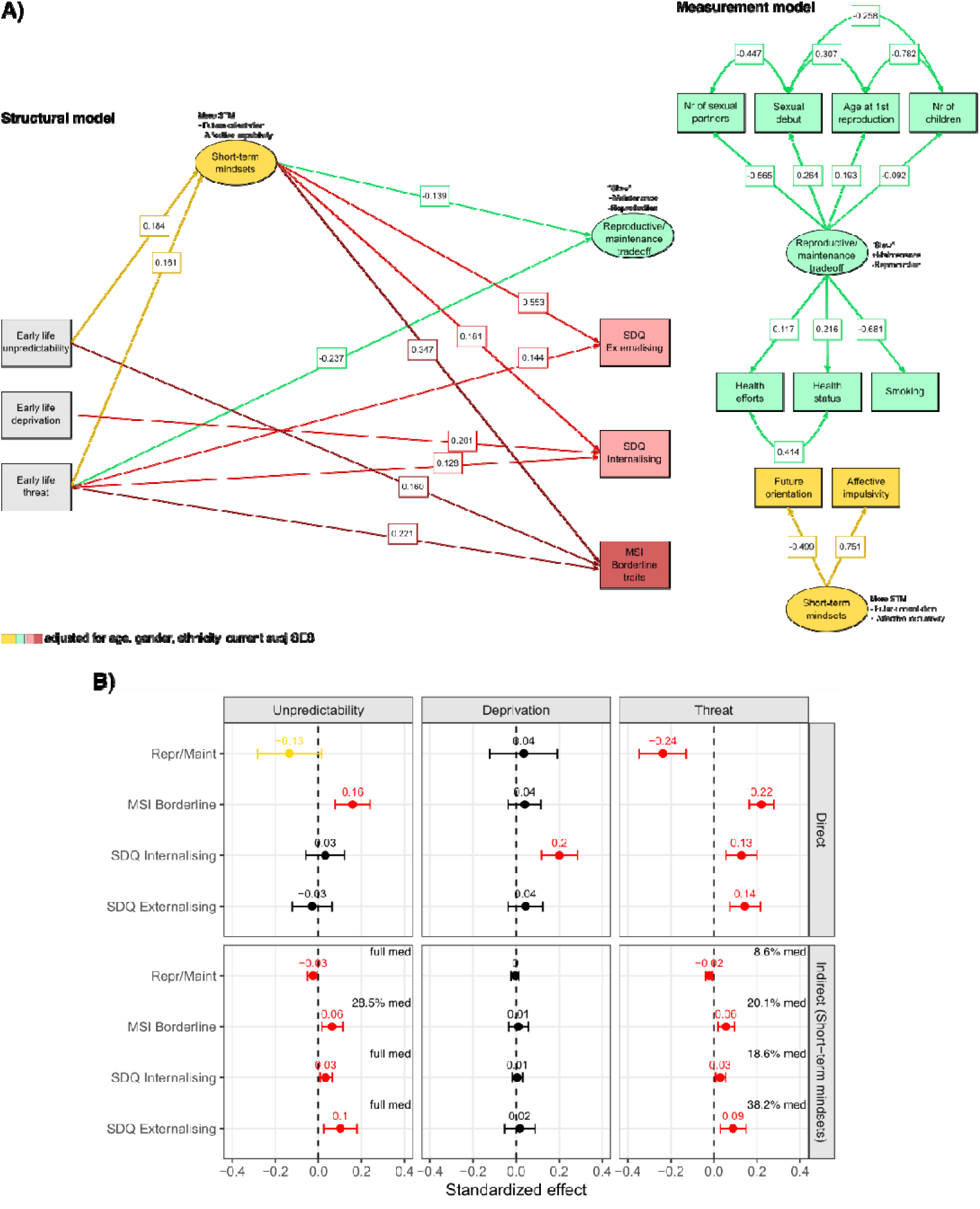
General unpredictability SEM. **A)** Simplified model diagram of the structural and measurement parts. Rectangles are manifest variables, ellipses are latent variables, double headed arrows are residual covariances, single headed arrows are regressions. Values are standardized coefficients. For visual clarity, not all paths are represented, only statistically significant regression paths and some of the residual covariances. Full list of model parameters can be found in Supplementary Table S4. **B)** Estimates of standardized direct and indirect effects of early life adversity dimensions on reproductive/maintenance and mental health. Error bars are Monte Carlo 95% confidence intervals. The proportion of total effect mediated is also shown in case of partial mediation (‘suppr’ and ‘full med’ indicate suppression and full mediation). Parameters significant at α level .05 are in red, Parameters significant at α level .1 are in yellow, parameters not significant are in black.

#### Measurement model

All indicators of the reproduction/maintenance effort trade-off loaded statistically significantly onto the latent, in directions matching theoretical expectations. The latent was correlated positively with sexual debut (β = .264, p < .001), and age at 1^st^ reproduction (β = .193, p < .001), but negatively with number of sexual partners (β = -.565, p < .001) and number of children (β = -.092, p = .010), indicative of later reproductive timing, less investment into mate seeking, and less reproductive output. The latent was correlated positively with both health status (β = .216, p < .001) and health efforts (β = .117, p = .007), but negatively with smoking (β = .681, p < .001), indicative of better subjective health and a greater effort of looking after one’s health. Thus, high scores on the latent seem to indicate that a participant is closer to following a ‘slow’ life history strategy, characterized by lesser reproductive, and greater somatic maintenance effort.

Both indicators of short-term mindsets also loaded significantly. High scores on the latent were characterized by greater affective impulsivity (β = .751, p < .001) and lower future orientation/more present orientation (β = -.499, p < .001), in other words, the adoption of short-term mindsets. The measurement model provides evidence for H1, suggesting that the covariance between our ‘biodemographic’ life history strategy indicators can be captured by a latent variable representing the hypothesized trade-off between reproductive effort and somatic maintenance effort.

#### Structural model

Short-term mindsets were predicted by both early life unpredictability (β = .184, p = .008) and threat (β = .161, p = .003), but not deprivation. In turn, short-term mindsets positively predicted all three mental health outcomes (Externalizing: β = .553, p < .001; Internalizing: β = .181, p < .001; Borderline: β = .347, p < .001), and negatively predicted the latent capturing greater maintenance and lesser reproductive effort (β = .139, p < .001). These indirect effects of unpredictability and threat were statistically significant at the .05 level for all outcomes according to Monte Carlo CIs (Figure 2B).

There were also direct effects of early life threat on all mental health outcomes (Externalizing: β = .144, p < .001; Internalizing: β = .128, p < .001; Borderline: β = .221, p < .001) and the reproduction/maintenance latent (β = -.237, p < .001). Early life unpredictability also had a direct effect on Borderline features (β = .160, p < .001) and small direct effect on the reproduction/maintenance latent significant at the .10 level (β = -.135, p = .082). Early life deprivation had a direct effect on internalizing (β = .201, p < .001).

The structural model provides evidence for H2. Greater exposure to early life adversity was associated with more mental health problems, and a shift towards ‘faster’ life history strategies, characterized by greater reproductive and lesser maintenance effort. Moreover, this was true especially for the dimension of threat. The model also provides evidence for H3. Greater exposure to early life adversity was associated with an indirect effect on mental health and the reproduction/maintenance trade-off through an increase in short-term mindsets, and this was especially pronounced for the dimension of unpredictability.

### Distinguishing short- and long-timescale unpredictability

Are these associations between early life unpredictability and mental health and lifestyle factors underlined by the same type of unpredictability? Or, similarly to deprivation and threat being partially dissociable features of harshness, can we meaningfully distinguish between different kinds of early unpredictability, with different consequences on development? We wished to explore this idea by splitting our unpredictability construct into an LT and ST component. The former being characterized by variability in environmental conditions on a longer timescale (months, years) leading to significantly changed contingencies, for example a residential move. The latter being characterised by variability on shorter timescales (hours, days, weeks) leading to ‘noisy’ variability around average contingencies that otherwise relatively stable on the long run, for example inconsistent parental discipline.

We classified eleven items of the QUIC as reflecting LT unpredictability (e.g., “There was a long period of time when I did not see one of my parents – e.g., military deployment, jail time, custody arrangements”), and twenty-one items as reflecting ST unpredictability (e.g., “One of my parents could go from calm to furious in an instant”). LT and ST individual scores were then computed by summing the responses for the relevant items. The complete list of items and their classification can be found in the Supplementary Table S1. The resulting subscales had adequate internal consistencies and inter-item correlations falling in the recommended range for scale unidimensionality (LT unpredictability: Cronbach α = .73 [.71, .76], average inter-item r = .21; ST unpredictability: Cronbach α = .88 [.87, .89], average inter-item r = .25).

It is important to note that while we report standard metrics of scale unidimensionality above, current theoretical frameworks caution against relying on factor analytic models to determine the psychometric validity of adversity measurements. The reason is that adversity dimensions should be defined not by how frequently certain experiences occur together, but by the similarity in their developmental mechanisms and their shared capacity to signal particular types of environmental adversity (Berman et al., 2022; Ellis et al., 2022; McLaughlin et al., 2021, 2023). For instance, while frequent parental employment changes and lack of family routines might co-occur frequently, we conceptualize the former as an indicator of LT environmental change and the latter as an indicator of ST environmental change, by virtue of their respective timescales and temporal signatures.

Despite the correlation *between items* of the subscales not being crucial in their definition for the reasons described above, the correlation *between the subscales* might be informative, since we would expect that if we capture different timescales of variability by our subscales, they might be statistically relatively more independent. To assess this, we created subscales with the same number of items as our original LT and ST subscales by randomly selecting items, with 10000 permutations, and calculated the correlation between each of these subscales. Remarkably, this revealed that the correlation between our a priori, theoretically derived split of items (r = .567) is lower than all correlations obtained from the permutations (Figure 3A).

**Figure 3.**
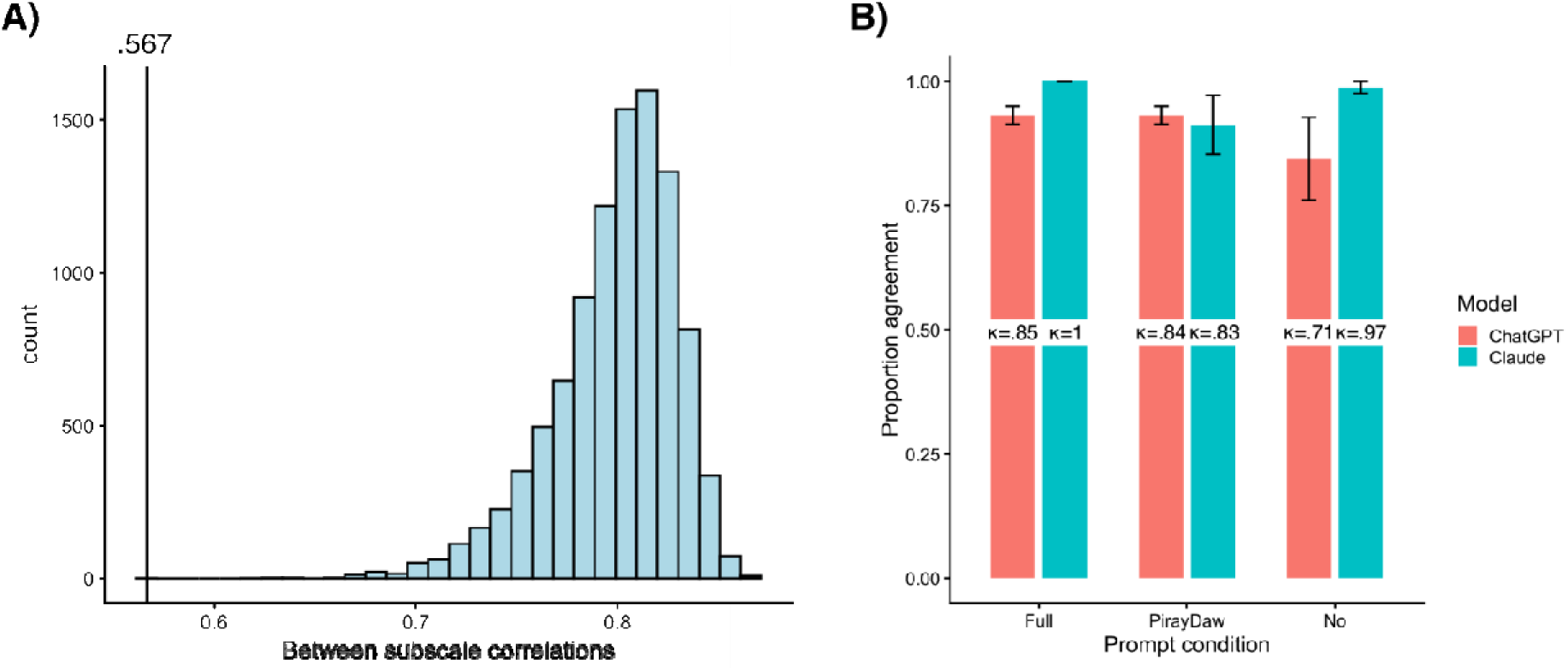
Validation of ST/LT subscales. **A)** Histogram showing the distribution of correlation coefficients between randomly created subscales with the number of items as the proposed ST and LT scales across 10000 permutations. The empirical correlation coefficient between our theory driven splits is also indicated by a vertical line. **B)** Classification agreement (mean ± standard error across 5 iterations with randomly permuted item order) and corresponding mean Cohen κ values between the LLMs (ChatGPT 5.3-mini in red, Claude Sonnet 4.6 in blue) and our theory driven item split in the different prompt conditions.

In order to assess the content validity of our theory-driven classification of items, we relied on large language models (LLM) as a more data-driven benchmark for content-based classification. We prompted GPT-5.3 and Claude Sonnet 4.6 to classify QUIC items into ST and LT based on the same criteria we used. It was provided with definitions of ST and LT unpredictability from either this manuscript, definitions of stochasticity and volatility from a recent RL study, or not provided with any further information (exact prompts are presented in Supplementary Text S1). The percentage of match between the LLMs’ classification and our own was high to perfect, independent of prompt condition, with Cohen’s κ ranging from 0.79 to 1 (Figure 3B). Given the acceptable distributions and reliability of the proposed scales, along with evidence for their relative independence and content validity, we used them in a subsequent model to estimate their shared and distinct associations with mental health and life history strategies.

### ST/LT SEM

#### Model fit and comparison

The ST/LT model presented with the following fit indices: χ^2^(57) = 303.11, p < .001; CFI = .939; RMSEA = .065 [.058, .073]; SRMR = .058. While fit indices are comparable (standardized residuals are presented in Supplementary Table S3), the ST/LT model had somewhat better 10-fold cross-validation performance in terms of the expected KL-divergence (E(KL-D) = 7.68 ± 0.76), compared to the main model (E(KL-D) = 8.04 ± 0.75). Overall, we deemed this model acceptable and retained it. A simplified diagram of model parameters is presented in Figure 4A, and the full table of fitted parameters is available in Supplementary Table S5.

**Figure 4.**
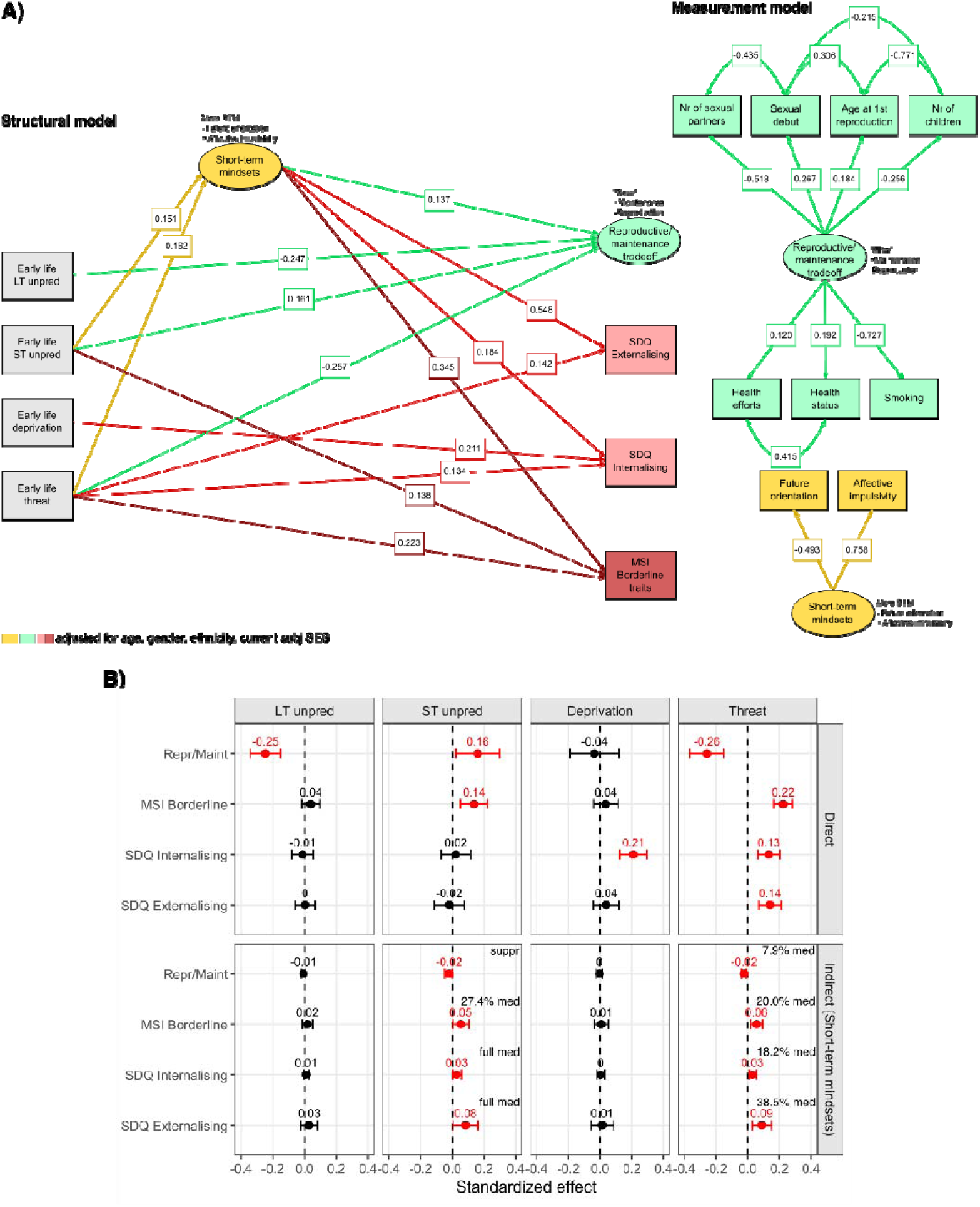
ST/LT SEM. **A)** Simplified model diagram of the structural and measurement parts. Rectangles are manifest variables, ellipses are latent variables, double headed arrows are residual covariances, single headed arrows are regressions. Values are standardized coefficients. For visual clarity, not all paths are represented, only statistically significant regression paths and some of the residual covariances. Full list of model parameters can be found in Supplementary Table S5. **B)** Estimates of standardized direct and indirect effects of early life adversity dimensions on reproductive/maintenance and mental health. Error bars are Monte Carlo 95% confidence intervals. The proportion of total effect mediated is also shown in case of partial mediation (‘suppr’ and ‘full med’ indicate suppression and full mediation). Parameters significant at α level .05 are in red, Parameters significant at α level .1 are in yellow, parameters not significant are in black.

#### Measurement model

For brevity, we do not detail the measurement model here again. All indicators of the reproduction/maintenance effort trade-off and short-term mindsets latents loaded statistically significantly, in directions matching theoretical expectations and the main model.

#### Structural model

Direct and indirect effects of both threat and deprivation remained unchanged from the main model. ST and LT unpredictability accounted for different components of the general unpredictability effects observed in the main model. LT unpredictability uniquely predicted reproduction-oriented life history strategies (β = -.247, p < .001). ST unpredictability accounted for all other effects. It had a direct effect on Borderline traits (β = .138, p = .002) and on short-term mindsets (β = .151, p = .036), which in turn continued to positively predict all three mental health outcomes (Externalizing: β = .549, p < .001; Internalizing: β = .184, p < .001; Borderline: β = .345, p < .001), and negatively predict the latent capturing greater maintenance and lesser reproductive effort (β = .137, p < .001). These indirect effects of ST unpredictability were all statistically significant (Figure 4B). Interestingly, ST unpredictability had a positive direct effect on the reproduction/maintenance latent (β = .161, p = .022), meaning it was associated with relatively greater maintenance over reproductive effort, contrary to other adversity dimensions.

To further test whether ST and LT unpredictability exert distinct effects across the model, we imposed equality constraints on all their corresponding structural paths. A scaled chi-square difference test indicated that these constraints significantly worsened model fit, Δχ²(5) = 26.32, p < .001. Consistent with this, a robust Wald test also rejected the joint equality of parameters, W(5) = 26.87, p < .001.

These results provide partial evidence for H4. While our model separating the early life ST and LT unpredictability showed only marginally better fit in terms of standard goodness of fit indices than our general unpredictability model, it did show clearly distinct effects of the two forms of early life uncertainty.

### Sensitivity analyses

#### Sexual debut only

We performed two sets of sensitivity analyses. First, rather than mixing direct LH traits (age at 1^st^ reproduction, fertility) and LH-related traits (sexual and health behaviours), we restricted our assessment of LH strategies to the sexual debut indicator only. Sexual debut has been proposed as a promising LH marker trait, that is relatively free from genetic confounds, responsive to environmental factors, and linked to other LH traits in a functionally coherent manner (Del Giudice, 2020, 2024).

This model fit acceptably, but worse than our main models: χ^2^(7) = 63.53, p < .001; CFI = .917; RMSEA = .096 [.075, .118]; SRMR = .037. All direct and indirect effects from the ST/LT model remained robust, with the exception of ST unpredictability’s direct and indirect effect on LH traits. Despite the model qualitatively recapturing the dissociation between ST and LT effects, neither the Chi square test, Δχ²(5) = 5.45, p = .364, nor the Wald test rejected the hypothesis of equality constraints, W(5) = 6.32, p = .277. In general, while the difference between ST and LT unpredictability coefficients weakened, the sensitivity analysis supports the validity of the main results.

#### Subjective adversity ratings

Second, rather than merely tallying the self-reported presence and frequency of individual adversity experiences, we replicated our model with adversity composite scores that were weighted by the subjective impact of each experience, according to participants. Specifically, we multiplied each item by a weight of 2 if its subjective impact was rated as ‘4-negative’, by a weight of 3 if its subjective impact was rated as ’5-very negative’, or left it with a weight of 1 otherwise. This scheme thus gives extra weight to experiences rated as having a more severe negative impact in our adversity sum scores, in line with recent calls to consider subjective severity in adversity research (Cohodes et al., 2023; Lacey & Minnis, 2020).

This model fit acceptably, but worse than our main models: χ^2^(64) = 334.55, p < .001; CFI = .937; RMSEA = .069 [.062, .077]; SRMR = .061. All direct and indirect effects from the ST/LT model remained robust, except for threat’s direct effect on internalizing problems. The model also qualitatively recaptured the dissociation between ST and LT unpredictability effects, and both the Chi square test, Δχ²(5) = 37.62, p < .001, and the Wald test were statistically significant, W(5) = 40.36, p < .001. In general, the sensitivity analysis supports the validity of the main results.

We also compared subjective severity ratings across adversity dimensions. Holm-corrected paired-samples t-tests indicated that ST unpredictability was rated as less severe than deprivation (p < .001), LT unpredictability (p = .008), and threat (p < .001). Deprivation was also rated as less severe than LT unpredictability (p = .002) and threat (p < .001), and LT unpredictability was rated as less severe than threat (p < .001). Descriptively, ST unpredictability received the lowest severity ratings (M = 2.44, SD = 0.61), followed by deprivation (M = 2.29, SD = 0.52) and LT unpredictability (M = 2.28, SD = 0.72), whereas threat was rated substantially higher (M = 1.15, SD = 0.70). Together, these results indicate a clear gradient in perceived severity, with threat evaluated as most impactful, followed by LT unpredictability and deprivation, and ST unpredictability rated as least severe. This also makes it clear that the specificity of indirect effects does not merely reflect the subjective severity of adversity dimensions, as both the lowest (ST unpredictability) and highest rated (threat) dimensions had mediated effects through short-term mindsets.

## Discussion

In this study, we tested four hypotheses regarding the association between dimensions of early life unpredictability, short-term mindsets, and mental health and LH outcomes. First, we expected that the covariation between indicators of reproductive and somatic maintenance effort will be accounted for by a latent variable corresponding with a trade-off between the two. We find supporting evidence for this, replicating earlier work (Baptista et al., 2023; Farkas et al., 2022; Lettinga et al., 2021; Mell et al., 2018). Across all models tested, a latent variable emerged, which captured better subjective health, more health efforts, less smoking, a later sexual debut, a smaller number of sexual partners, lower fertility, and later age at 1^st^ reproduction. In other words, greater maintenance over reproductive effort. Smoking, sexual debut, and number of sexual partners proved to be strongest loading indicators of the latent. Indeed, sexual debut has been recently proposed as a particularly suitable marker of LH strategies, given its coherent association with other LH indicators and the relatively large environmental contributions to its phenotypic variance (Del Giudice, 2020, 2024). The small contribution of fertility to our latent is also not surprising, as in a Western, post demographic transition context, fertility has been somewhat decoupled from reproductive behaviour due to cultural factors (Lawson & Borgerhoff Mulder, 2016), and accordingly, this indicator has been found to be weak or nonsignificant in earlier SEM based LH research (Lettinga et al., 2021; Mell et al., 2018). It is important to note, that in line with recent calls in the field (Međedović, 2020; Sear, 2020), we measured a functional LH trade-off using both psychobehavioural and the biodemographic indices, rather than using psychometric measures or reproductive indicators of a “fast” LH strategy only.

Our second hypothesis centred on the effects of early life adversity dimensions on LH and mental health outcomes, and their potential mediation through the induction of short-term mindsets. We expected that early life adversity would be associated with a shift towards greater reproductive over maintenance effort, and worse mental health, and that these effects would be at least partially mediated by shorter-term mindsets. We expected relatively stronger direct effects of threat, given its robust association with socioemotional development and psychopathology in previous studies (Ellis et al., 2022; A. H. Lee et al., 2024), and its close theoretical link to extrinsic mortality/morbidity (Ellis et al., 2009); as well as relatively stronger indirect effects of early life unpredictability, given its proposed role in regulating unpredictability schemas and short-term mindsets (Cabeza de Baca et al., 2016; Deitzer et al., 2025; Ellis et al., 2022; Szepsenwol et al., 2017). We also found confirmatory evidence for these hypotheses. With respect to effects independent of short-term mindsets, threat had significant direct links to all three mental health outcomes, and with the reproduction/maintenance latent in the expected direction. Deprivation was only directly associated with internalizing problems, and unpredictability with Borderline traits. A recent meta-analysis also found that while effect sizes of threat’s association with internalizing, externalizing, and PTSD symptoms were consistently larger than deprivation’s, deprivation’s association was strongest for internalizing (Lee et al., 2024). This is likely driven to a large extent by the inclusion of emotional neglect in this dimension, which is strongly linked to internalizing problems (Hildyard & Wolfe, 2002), although exact mechanisms are still not clear, with proposed roles for alexithymia (Brown et al., 2016), emotional dysregulation (Shin et al., 2025; Zhang et al., 2024), and perceived control (Bolger & Patterson, 2001). Given the central role of mood and interpersonal instability in borderline personality disorder, its direct links with early life unpredictability are not surprising (Gunderson et al., 2018). The role of early life adversity, and the ensuing insecure attachment styles, and mentalizing difficulties are thought to play central roles in the aetiology of the condition (Fonagy & Bateman, 2008; Levy, 2005). These results replicate earlier findings and indicate broad effects of early life threat on mental health and LH strategies, that are independent of short-term mindsets. This pattern can be interpreted in light of psychosocial acceleration theory and its later elaborations, in which energetic stress constrains a first ‘tier’ of LH responses that delay reproductive development, whereas psychosocial stress accelerates later life reproductive and health behaviour in a second ‘tier’ (Belsky et al., 1991; Carrera et al., 2026; Ellis et al., 2024).

There was also evidence that short-term mindsets mediated adversity – outcome associations. They were strongly associated with all three mental health outcomes, as well as with a reproduction-oriented LH strategy. We observed partial mediation of early life threat’s effects on all outcomes, and complete mediation of early life unpredictability’s effects on internalizing, externalizing, and reproduction-oriented LH strategies. These effects are entirely in line with evolutionary-developmental models of psychopathology (Del Giudice, 2018; Ellis et al., 2022) and previous results robustly linking early life harshness and unpredictability to “fast” LH strategies and their psychological mediators, and increased risk for externalizing psychopathology and risk-taking (Doom et al., 2016; Farkas & Jacquet, 2024; Hartman et al., 2018; Martinez et al., 2022; Mell et al., 2018; Simpson et al., 2012; Szepsenwol et al., 2017; Usacheva et al., 2022). They also go beyond earlier studies by capturing both a cognitive (future orientation) and affective/behavioural (affective impulsivity) component of the short-term mindset in a more comprehensive manner. Previous studies tended to focus on a single component process, leading to a fragmented and incomplete understanding of the construct, an issue recently highlighted by van Gelder et al. (2025) and van Gelder & Frankenhuis (2025). Our results on the other hand, indicate that cognitive and affective/behavioural facets of short-term mindsets covary systematically, follow unpredictable early environments, and mediate their effects on mental health as well as reproductive and health behaviours.

We wanted to examine these early life unpredictability associations in more detail, given the uncertainty in the conceptualization and measurement of the construct. We explored an approach, inspired by RL, in which environmental unpredictability is subdivided into more stochastic, short-term (ST), and more volatile, long-term (LT) forms. To this end, we created two subscales from the existing QUIC questionnaire, assessed their content validity and reliability, and investigated their shared and distinct associations with short-term mindsets and outcomes. This analysis revealed an interesting dissociation, in which ST and LT unpredictability accounted for different parts of the general unpredictability effects uncovered in our earlier analysis. ST unpredictability seemed responsible for the direct effect on Borderline traits, as well as for all indirect effects through short-term mindsets. LT unpredictability was uniquely associated with a shift toward reproductive over maintenance effort instead. Interestingly, ST unpredictability had a smaller direct effect in the opposite direction, favouring maintenance effort. While few previous studies focused on decomposing early life unpredictability, there are some findings that can inform the interpretation of these results. An early study focusing on specific sources of environmental instability has shown that longer-term, structural changes – especially parental transitions – are particularly strong predictors of reproductive behaviour, risk taking, and externalising outcomes (Hartman et al., 2018), aligning with the association we observed between LT unpredictability and reproductive effort. Similarly, studies distinguishing between forms of adversity often find that more enduring or event-like disruptions (e.g., negative life events or income fluctuations over time) are more consistently linked to externalising behaviour, whereas more proximal, chaotic or noisy environments show different or weaker patterns of association with behavioural outcomes (Chang et al., 2021; Z. Li & Belsky, 2022). A recent study further suggests that economic unpredictability itself might be decomposed into a component marked by frequent, unpredictable hardship, and another marked by infrequent but abrupt hardship, somewhat mirroring our distinction, with a weighted composite unpredictability index predicting self-regulation challenges over and above mean economic hardship (DeJoseph et al., 2025).

Our findings are also consistent with emerging evidence that more explicitly considers the timescale of unpredictability (Davis & Glynn, 2024). In a longitudinal cohort study, separating ST (more stochastic) and LT (more volatile) unpredictability improved the prediction of both internalising and externalising behaviours, with LT unpredictability showing particularly robust associations with externalising outcomes (Farkas et al., 2024). Notably, this study also observed an interaction between these dimensions, suggesting that developmental outcomes may be most pronounced under low ST and high LT uncertainty, a pattern similar to the countervailing effects we observed on reproductive and health behaviour. In addition, the study also found a mediation effect of reproductive strategy uniquely for the LT component. Finally, work on early sensory environments indicates that the statistical structure (their autocorrelation) of inputs has measurable effects on attentional and neural processing from infancy (Werchan et al., 2022). Our distinction also maps onto the difference between the ‘ancestral cue’ and ‘statistical learning’ sources of unpredictability, the former referring to individual events (e.g., parental separations) that natural selection has shaped the human brain to treat as reliable indicators of unpredictability, the latter referring to the brain’s general monitoring of environmental states in a general sense (Young et al., 2020). One possibility is that stochastic, ST variability has a stronger effect on the adoption of unpredictability schemas and short-term mindsets through engaging statistical learning processes, whereas volatile, LT changes might be picked up as ancestral cues and more calibrate behavioural components of LH strategies through different pathways, such as physiological stress reactivity (Li et al., 2023). Taken together, these findings suggest that timescale is a theoretically and empirically useful dimension for refining the study of unpredictability.

A number of limitations should also be noted. First, the study is cross-sectional and relies on adults’ retrospective reports of childhood experiences, which limits causal inference, given the often modest correspondence between retrospectively and prospectively assessed adverse experiences (Baldwin et al., 2019; Coleman et al., 2026; Reuben et al., 2016). Although future work should prioritize validation using objective and longitudinal designs, it is worth noting that concordance between subjective and objective measures is not uniformly low (Naicker et al., 2021; Patten et al., 2015), and subjective appraisals of adversity have, in some cases, been shown to be equal or stronger predictors of functional outcomes than objective indicators (Danese & Widom, 2020; Francis et al., 2023; Rivenbark et al., 2020). Second, several constructs were necessarily approximated with a degree of imprecision. Our reproductive-maintenance latent captures a biodemographic trade-off only imperfectly, and the ST/LT split was derived from theory-guided item classification within an existing unpredictability measure rather than from a purpose-built instrument. Relatedly, some measures were modified from their original form, including restricting all adversity reports to experiences before age 12 and adding subjective impact ratings. Finally, although the sample was large and approximately representative of the UK adult population, generalizability beyond this cultural context remains uncertain, and the fit of the SEMs, while acceptable, was not uniformly strong enough to rule out alternative model specifications.

Despite these limitations, the present findings support a dimensional and evolutionary-developmental account of childhood adversity in which threat, deprivation, and unpredictability show partially dissociable links with adult mental health and life history-related outcomes, and in which short-term mindsets emerge as one plausible mediating pathway. Furthermore, they suggest that unpredictability itself is not a unitary construct: shorter-timescale, more stochastic experiences were more closely tied to short-term mindsets and psychopathology, whereas longer-timescale, more volatile experiences were more specifically linked to a shift toward reproductive over maintenance effort.

## Author Contributions

Bence C. Farkas : Conceptualization, Methodology, Software, Validation, Formal analysis, Investigation, Data curation, Writing - Original Draft, Writing - Review & Editing, Visualization

Valentin Wyart : Conceptualization, Methodology, Resources, Writing - Review & Editing, Supervision, Project administration, Funding acquisition

Pierre O. Jacquet : Conceptualization, Methodology, Resources, Writing - Review & Editing, Supervision, Project administration, Funding acquisition

