## Supplementary Materials for "Early life adversity is associated with mental health and life histories through short-term mindsets"

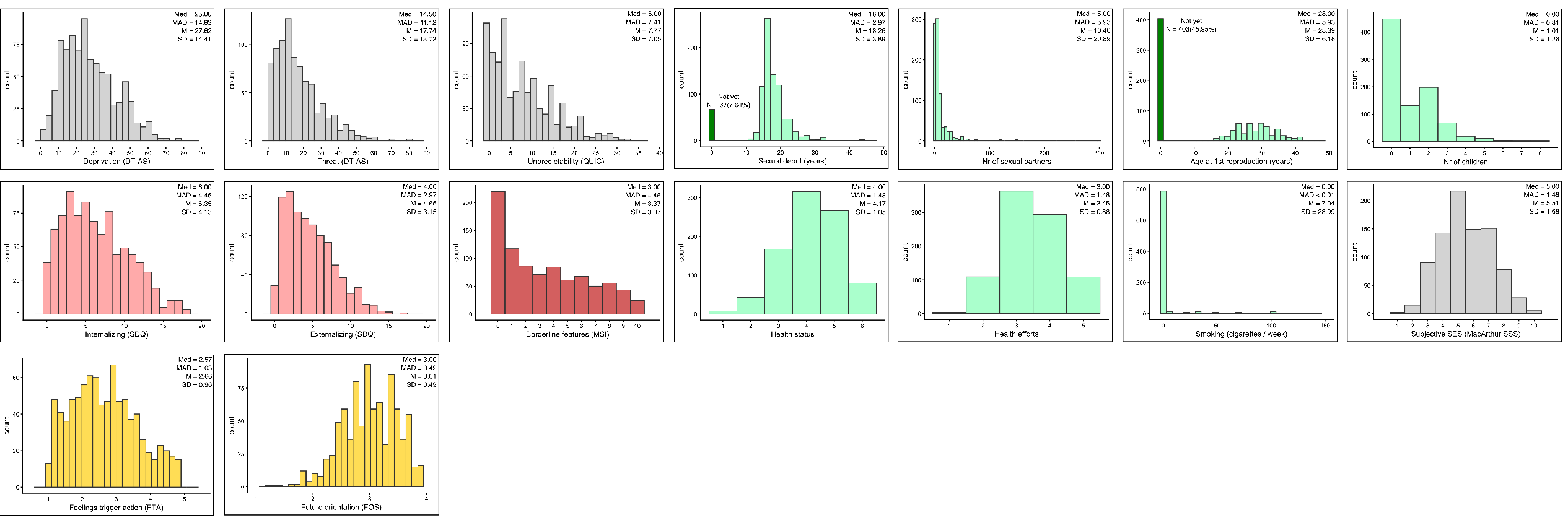

**Supplementary Figure S1. Distributions and descriptive statistics of study variables.**

| **QUIC Nr.** | **Item wording** | **Dimension** |
| --- | --- | --- |
| 1. | I had a set morning routine on school days (i.e., I usually did the same thing each day to get ready). (R) | ST unpred |
| 2. | My parents kept track of what I ate (e.g., made sure that I didn’t skip meals or tried to make sure I ate healthy food). (R) | ST unpred |
| 3. | My family ate a meal together most days. (R) | ST unpred |
| 4. | My parents tried to make sure I got a good night’s sleep (e.g., I had a regular bedtime, my parents checked to make sure I went to sleep). (R) | ST unpred |
| 5. | I had a bedtime routine (e.g., my parents tucked me in, my parents read me a book, I took a bath). (R) | ST unpred |
| 6. | In my afterschool or free time hours at least one of my parents knew what I was doing. (R) | ST unpred |
| 7. | At least one of my parents regularly checked that I did my homework. (R) | ST unpred |
| 8. | At least one of my parents regularly kept track of my school progress. (R) | ST unpred |
| 9. | At least one parent made time each day to see how I was doing. (R) | ST unpred |
| 10. | My parents were often late to pick me up (e.g. from school, aftercare or sports) | ST unpred |
| 11. | I usually knew when my parents were going to be home. (R) | ST unpred |
| 12. | At least one of my parents had punishments that were unpredictable | ST unpred |
| 13. | I often wondered whether or not one of my parents would come home at the end of the day | ST unpred |
| 14. | My family planned activities to do together. (R) | ST unpred |
| 15. | At least one of my parents would plan something for the family, but then not follow through with the plan | ST unpred |
| 16. | My family had holiday traditions that we did every year (e.g., cooking a special food at a particular time of year/decorate the house the same way). (R) | LT unpred |
| 17. | At least one of my parents was disorganized | ST unpred |
| 18. | At least one of my parents was unpredictable | ST unpred |
| 19. | For at least one of my parents, when they were upset, I did not know how they would act | ST unpred |
| 20. | One of my parents could go from calm to furious in an instant | ST unpred |
| 21. | One of my parents could go from calm to stressed or nervous in an instant | ST unpred |
| 22. | There was a long period of time when I didn’t see one of my parents (e.g. military deployment, jail time, custody arrangements) | LT unpred |
| 23. | I experienced changes in my custody arrangement | LT unpred |
| 24. | At least one of my parents changed jobs frequently | LT unpred |
| 25. | There were times when one of my parents was unemployed and couldn’t find a job even though he/she wanted one | LT unpred |
| 26. | My parents had a stable relationship with each other. (R) | LT unpred |
| 27. | My parents got divorced | LT unpred |
| 28. | At least one of my parents had many romantic partners | LT unpred |
| 29. | There were often people coming and going in my house that I did not expect to be there | ST unpred |
| 30. | I moved frequently | LT unpred |
| 31. | I changed schools frequently | LT unpred |
| 32. | I changed schools mid-year | LT unpred |
| 33. | I lived in a clean house. (R) | - |
| 34. | I lived in a cluttered house (e.g., piles of stuff everywhere) | - |
| 35. | In my house things I needed were often misplaced so that I could not find them | - |
| 36. | There was a period of time when I often worried that I was not going to have enough food to eat | - |
| 37. | There was a period of time when I often worried that my family would not have enough money to pay for necessities like clothing or bills | - |
| 38. | There was a period of time when I did not feel safe in my home | - |

**Supplementary Table S1. QUIC items and their classification into ST and LT unpredictability.**

**Supplementary Table S2. Residual correlations of the general unpredictability SEM.** Correlations above r = .10 are highlighted in bold.

|  | Sexual debut | Nr of sexual partners | Age at 1^st^ repr. | Nr of children | Health status | Health efforts | Smoking | Future orientation | Aff. impulsivity | Externalizing | Internalizing | Borderline |
| --- | --- | --- | --- | --- | --- | --- | --- | --- | --- | --- | --- | --- |
| Sexual debut | - |  |  |  |  |  |  |  |  |  |  |  |
| Nr of sexual partners | < .01 | - |  |  |  |  |  |  |  |  |  |  |
| Age at 1^st^ repr | < .01 | .03 | - |  |  |  |  |  |  |  |  |  |
| Nr of children | < .01 | .01 | < .01 | - |  |  |  |  |  |  |  |  |
| Health status | - .07 | .10 | - .09 | .10 | - |  |  |  |  |  |  |  |
| Health efforts | - .06 | .02 | - .02 | - .03 | < .01 | - |  |  |  |  |  |  |
| Smoking | .02 | < .01 | - .02 | < .01 | - .03 | < .01 | - |  |  |  |  |  |
| Future orientation | **.12** | - .10 | .03 | - .02 | .03 | **.17** | - .09 | - |  |  |  |  |
| Aff. impulsivity | - .03 | - .06 | **- .12** | **.12** | **- .16** | **- .17** | - .05 | < .01 | - |  |  |  |
| Externalizing | .02 | .02 | .07 | - .02 | < .01 | < .01 | - .01 | - .03 | - .03 | - |  |  |
| Internalizing | **.18** | - .02 | .08 | - .07 | < .01 | < .01 | .05 | .12 | .10 | < .01 | - |  |
| Borderline | .01 | .03 | .05 | - .05 | < .01 | < .01 | - .01 | .07 | .06 | < .01 | < .01 | - |

|  | Sexual debut | Nr of sexual partners | Age at 1^st^ repr. | Nr of children | Health status | Health efforts | Smoking | Future orientation | Aff. impulsivity | Internalizing | Externalizing | Borderline |
| --- | --- | --- | --- | --- | --- | --- | --- | --- | --- | --- | --- | --- |
| Sexual debut | - |  |  |  |  |  |  |  |  |  |  |  |
| Nr of sexual partners | < .01 | - |  |  |  |  |  |  |  |  |  |  |
| Age at 1^st^ repr | < .01 | .03 | - |  |  |  |  |  |  |  |  |  |
| Nr of children | < .01 | - .08 | < .01 | - |  |  |  |  |  |  |  |  |
| Health status | - .07 | .07 | - .09 | .13 | - |  |  |  |  |  |  |  |
| Health efforts | - .06 | .02 | - .01 | - .01 | < .01 | - |  |  |  |  |  |  |
| Smoking | .05 | < .01 | - .01 | .04 | - .03 | < .01 | - |  |  |  |  |  |
| Future orientation | **.13** | - .10 | .03 | - .01 | .03 | **.17** | - .09 | - |  |  |  |  |
| Aff. impulsivity | - .03 | - .06 | **- .12** | .10 | **- .16** | **- .17** | - .06 | < .01 | - |  |  |  |
| Internalizing | .02 | .02 | .07 | - .02 | < .01 | < .01 | - .01 | - .04 | - .03 | - |  |  |
| Externalizing | **.18** | - .02 | .08 | - .05 | < .01 | < .01 | .05 | .12 | .10 | < .01 | - |  |
| Borderline | .01 | .03 | .05 | - .06 | < .01 | < .01 | - .01 | .07 | .05 | < .01 | < .01 | - |

**Supplementary Table S3. Residual correlations of the ST/LT SEM.** Correlations above r = .10 are highlighted in bold.

|  | Term | b (SE b) | 95% CI | β | z | p |
| --- | --- | --- | --- | --- | --- | --- |
| Latent variables | **Repr/Maint =~ Sexual debut** | 0.252 (0.056) | [0.142, 0.362] | 0.264 | 4.493 | **<.001** |
|  | **Repr/Maint =~**  **Nr sexual partners** | -0.518 (0.034) | [-0.585, -0.451] | -0.565 | -15.248 | **<.001** |
|  | **Repr/Maint =~**  **Age 1^st^ reproduction** | 0.208 (0.025) | [0.159, 0.257] | 0.193 | 8.175 | **<.001** |
|  | **Repr/Maint =~ Nr children** | -0.101 (0.039) | [-0.177, -0.025] | -0.092 | -2.593 | **.010** |
|  | **Repr/Maint =~ Health status** | 0.205 (0.023) | [0.160, 0.250] | 0.216 | 8.841 | **<.001** |
|  | **Repr/Maint =~ Health efforts** | 0.111 (0.042) | [0.029, 0.193] | 0.117 | 2.674 | **.007** |
|  | **Repr/Maint =~ Smoking** | -0.641 (0.045) | [-0.729, -0.553] | -0.681 | -14.332 | **<.001** |
|  | **Short-term mds =~ Future ori** | 0.233 (0.018) | [0.198, 0.268] | 0.499 | 12.684 | **<.001** |
|  | **Short-term mds =~**  **Affective imp** | -0.675 (0.046) | [-0.765, -0.585] | -0.751 | -14.619 | **<.001** |
| Regressions | **Short-term mds ~**  **Early life unp** | -0.028 (0.010) | [-0.048, -0.008] | -0.184 | -2.640 | **.008** |
|  | **Short-term mds ~**  **Early life threat** | -0.012 (0.004) | [-0.020, -0.004] | -0.161 | -2.951 | **.003** |
|  | Short-term mds ~  Early life depr | -0.002 (0.005) | [-0.012, 0.008] | -0.030 | -0.467 | .640 |
|  | Repr/Maint ~  Early life unp | -0.021 (0.012) | [-0.045, 0.003] | -0.135 | -1.741 | .082 |
|  | **Repr/Maint ~**  **Early life threat** | -0.019 (0.005) | [-0.029, -0.009] | -0.237 | -4.121 | **<.001** |
|  | Repr/Maint ~  Early life depr | 0.003 (0.006) | [-0.009, 0.015] | 0.036 | 0.438 | .662 |
|  | **Repr/Maint ~**  **Short-term mds** | 0.142 (0.029) | [0.085, 0.199] | 0.139 | 4.828 | **<.001** |
|  | Externalizing ~  Early life unp | -0.013 (0.021) | [-0.054, 0.028] | -0.029 | -0.624 | .533 |
|  | **Externalizing ~**  **Early life threat** | 0.033 (0.008) | [0.017, 0.049] | 0.144 | 3.999 | **<.001** |
|  | Externalizing ~  Early life depr | 0.010 (0.009) | [-0.008, 0.028] | 0.044 | 1.066 | .287 |
|  | **Externalizing ~**  **Short-term mds** | -1.644 (0.110) | [-1.860, -1.428] | -0.553 | -14.990 | **<.001** |
|  | Internalizing ~  Early life unp | 0.019 (0.027) | [-0.034, 0.072] | 0.032 | 0.700 | .484 |
|  | **Internalizing ~**  **Early life threat** | 0.038 (0.011) | [0.016, 0.060] | 0.128 | 3.510 | **<.001** |
|  | **Internalizing ~**  **Early life depr** | 0.058 (0.012) | [0.034, 0.082] | 0.201 | 4.673 | **<.001** |
|  | **Internalizing ~**  **Short-term mds** | -0.706 (0.136) | [-0.973, -0.439] | -0.181 | -5.184 | **<.001** |
|  | **Borderline ~**  **Early life unp** | 0.069 (0.018) | [0.034, 0.104] | 0.160 | 3.775 | **<.001** |
|  | **Borderline ~**  **Early life threat** | 0.049 (0.007) | [0.035, 0.063] | 0.221 | 7.459 | **<.001** |
|  | Borderline ~  Early life depr | 0.008 (0.008) | [-0.008, 0.024] | 0.040 | 1.026 | .305 |
|  | **Borderline ~**  **Short-term mds** | -1.001 (0.100) | [-1.197, -0.805] | -0.347 | -10.046 | **<.001** |
| Residual covariances | **Health status ~~ Health efforts** | 0.403 (0.026) | [0.352, 0.454] | 0.414 | 15.506 | **<.001** |
|  | **Sexual debut ~~**  **Nr sexual partners** | -0.351 (0.041) | [-0.431, -0.271] | -0.447 | -8.645 | **<.001** |
|  | **Sexual debut ~~**  **Age 1^st^ reproduction** | 0.290 (0.037) | [0.218, 0.362] | 0.307 | 7.760 | **<.001** |
|  | **Sexual debut ~~**  **Nr children** | -0.248 (0.041) | [-0.328, -0.168] | -0.258 | -6.024 | **<.001** |
|  | **Age 1^st^ reproduction ~~**  **Nr children** | -0.760 (0.018) | [-0.795, -0.725] | -0.782 | -41.845 | **<.001** |
|  | **Health status ~~ Internalizing** | -1.177 (0.115) | [-1.402, -0.952] | -0.353 | -10.222 | **<.001** |
|  | **Health status ~~ Externalizing** | -0.569 (0.095) | [-0.755, -0.383] | -0.264 | -5.999 | **<.001** |
|  | **Health status ~~ Borderline** | -0.638 (0.085) | [-0.805, -0.471] | -0.294 | -7.466 | **<.001** |
|  | **Health efforts ~~ Internalizing** | -0.703 (0.115) | [-0.928, -0.478] | -0.208 | -6.132 | **<.001** |
|  | **Health efforts ~~ Externalizing** | -0.401 (0.095) | [-0.587, -0.215] | -0.183 | -4.216 | **<.001** |
|  | **Health efforts ~~ Borderline** | -0.320 (0.082) | [-0.481, -0.159] | -0.145 | -3.912 | **<.001** |
|  | Repr/Maint ~~  Externalizing | 0.034 (0.052) | [-0.068, 0.136] | 0.016 | 0.654 | .513 |
|  | **Repr/Maint ~~**  **Internalizing** | 0.482 (0.098) | [0.290, 0.674] | 0.141 | 4.914 | **<.001** |
|  | Repr/Maint ~~  Borderline | -0.067 (0.041) | [-0.147, 0.013] | -0.030 | -1.607 | .108 |
|  | **Externalizing ~~**  **Internalizing** | 2.744 (0.299) | [2.158, 3.330] | 0.366 | 9.190 | **<.001** |
|  | **Externalizing ~~**  **Borderline** | 1.512 (0.213) | [1.094, 1.930] | 0.309 | 7.101 | **<.001** |
|  | **Internalizing ~~**  **Borderline** | 3.715 (0.308) | [3.111, 4.319] | 0.491 | 12.059 | **<.001** |

**Supplementary Table S4. Parameter estimates of the General unpredictability SEM.** Regressions of adjustment variables, intercepts, thresholds and variances are not included for readability (altogether they would add 89 additional parameters), but the full model is available in the supplementary .RData file. Statistically significant terms are highlighted in bold.

|  | Term | b (SE b) | 95% CI | β | z | p |
| --- | --- | --- | --- | --- | --- | --- |
| Latent variables | **Repr/Maint =~ Sexual debut** | 0.254 (0.052) | [0.152, 0.356] | 0.267 | 4.920 | **<.001** |
|  | **Repr/Maint =~**  **Nr sexual partners** | -0.467 (0.027) | [-0.520, -0.414] | -0.518 | -17.566 | **<.001** |
|  | **Repr/Maint =~**  **Age 1^st^ reproduction** | 0.198 (0.025) | [0.149, 0.247] | 0.184 | 8.003 | **<.001** |
|  | **Repr/Maint =~ Nr children** | -0.278 (0.032) | [-0.341, -0.215] | -0.256 | -8.685 | **<.001** |
|  | **Repr/Maint =~ Health status** | 0.180 (0.023) | [0.135, 0.225] | 0.192 | 8.017 | **<.001** |
|  | **Repr/Maint =~ Health efforts** | 0.113 (0.039) | [0.037, 0.189] | 0.120 | 2.885 | **.004** |
|  | **Repr/Maint =~ Smoking** | -0.679 (0.042) | [-0.761, -0.597] | -0.727 | -16.245 | **<.001** |
|  | **Short-term mds =~ Future ori** | 0.230 (0.018) | [0.195, 0.265] | 0.493 | 12.516 | **<.001** |
|  | **Short-term mds =~**  **Affective imp** | -0.683 (0.047) | [-0.775, -0.591] | -0.758 | -14.600 | **<.001** |
| Regressions | **Short-term mds ~**  **Early life stoch** | -0.034 (0.016) | [-0.065, -0.003] | -0.151 | -2.093 | **.036** |
|  | Short-term mds ~  Early life volat | -0.027 (0.024) | [-0.074, 0.020] | -0.052 | -1.083 | .279 |
|  | **Short-term mds ~**  **Early life threat** | -0.012 (0.004) | [-0.020, -0.004] | -0.162 | -2.989 | **.003** |
|  | Short-term mds ~  Early life depr | -0.002 (0.005) | [-0.012, 0.008] | -0.024 | -0.372 | .710 |
|  | **Repr/Maint ~**  **Early life stoch** | 0.038 (0.016) | [0.007, 0.069] | 0.161 | 2.283 | **.022** |
|  | **Repr/Maint ~**  **Early life volat** | -0.129 (0.026) | [-0.180, -0.078] | -0.247 | -4.879 | **<.001** |
|  | **Repr/Maint ~**  **Early life threat** | -0.020 (0.004) | [-0.028, -0.012] | -0.257 | -4.551 | **<.001** |
|  | Repr/Maint ~  Early life depr | -0.003 (0.006) | [-0.015, 0.009] | -0.036 | -0.453 | .650 |
|  | **Repr/Maint ~**  **Short-term mds** | 0.141 (0.031) | [0.080, 0.202] | 0.137 | 4.604 | **<.001** |
|  | Externalizing ~  Early life stoch | -0.013 (0.033) | [-0.078, 0.052] | -0.020 | -0.409 | .682 |
|  | Externalizing ~  Early life volat | 0.005 (0.048) | [-0.089, 0.099] | 0.003 | 0.102 | .919 |
|  | **Externalizing ~**  **Early life threat** | 0.033 (0.008) | [0.017, 0.049] | 0.142 | 3.955 | **<.001** |
|  | Externalizing ~  Early life depr | 0.009 (0.009) | [-0.009, 0.027] | 0.039 | 0.944 | .345 |
|  | **Externalizing ~**  **Short-term mds** | -1.632 (0.110) | [-1.848, -1.416] | -0.549 | -14.850 | **<.001** |
|  | Internalizing ~  Early life stoch | 0.020 (0.043) | [-0.064, 0.104] | 0.022 | 0.459 | .646 |
|  | Internalizing ~  Early life volat | -0.025 (0.068) | [-0.158, 0.108] | -0.012 | -0.362 | .717 |
|  | **Internalizing ~**  **Early life threat** | 0.040 (0.011) | [0.018, 0.062] | 0.134 | 3.668 | **<.001** |
|  | **Internalizing ~**  **Early life depr** | 0.060 (0.013) | [0.035, 0.085] | 0.211 | 4.770 | **<.001** |
|  | **Internalizing ~**  **Short-term mds** | -0.716 (0.136) | [-0.983, -0.449] | -0.184 | -5.252 | **<.001** |
|  | **Borderline ~**  **Early life stoch** | 0.091 (0.029) | [0.034, 0.148] | 0.138 | 3.097 | **.002** |
|  | Borderline ~  Early life volat | 0.057 (0.042) | [-0.025, 0.139] | 0.039 | 1.359 | .174 |
|  | **Borderline ~**  **Early life threat** | 0.050 (0.007) | [0.036, 0.064] | 0.223 | 7.584 | **<.001** |
|  | Borderline ~  Early life depr | 0.008 (0.008) | [-0.008, 0.024] | 0.036 | 0.910 | .363 |
|  | **Borderline ~**  **Short-term mds** | -0.998 (0.100) | [-1.194, -0.802] | -0.345 | -10.005 | **<.001** |
| Residual covariances | **Health status ~~ Health efforts** | 0.405 (0.026) | [0.354, 0.456] | 0.415 | 15.729 | **<.001** |
|  | **Sexual debut ~~**  **Nr sexual partners** | -0.350 (0.033) | [-0.415, -0.285] | -0.435 | -10.774 | **<.001** |
|  | **Sexual debut ~~**  **Age 1^st^ reproduction** | -0.725 (0.019) | [-0.762, -0.688] | -0.771 | -37.512 | **<.001** |
|  | **Sexual debut ~~**  **Nr children** | 2.742 (0.298) | [2.158, 3.326] | 0.364 | 9.204 | **<.001** |
|  | **Age 1^st^ reproduction ~~**  **Nr children** | 1.517 (0.213) | [1.100, 1.934] | 0.309 | 7.106 | **<.001** |
|  | **Health status ~~ Internalizing** | 3.728 (0.308) | [3.124, 4.332] | 0.493 | 12.096 | **<.001** |
|  | **Health status ~~ Externalizing** | 0.254 (0.052) | [0.152, 0.356] | 0.267 | 4.920 | **<.001** |
|  | **Health status ~~ Borderline** | -0.467 (0.027) | [-0.520, -0.414] | -0.518 | -17.566 | **<.001** |
|  | **Health efforts ~~ Internalizing** | 0.198 (0.025) | [0.149, 0.247] | 0.184 | 8.003 | **<.001** |
|  | **Health efforts ~~ Externalizing** | -0.278 (0.032) | [-0.341, -0.215] | -0.256 | -8.685 | **<.001** |
|  | **Health efforts ~~ Borderline** | 0.180 (0.023) | [0.135, 0.225] | 0.192 | 8.017 | **<.001** |
|  | **Repr/Maint ~~**  **Externalizing** | 0.113 (0.039) | [0.037, 0.189] | 0.120 | 2.885 | **.004** |
|  | **Repr/Maint ~~**  **Internalizing** | -0.679 (0.042) | [-0.761, -0.597] | -0.727 | -16.245 | **<.001** |
|  | **Repr/Maint ~~**  **Borderline** | 0.230 (0.018) | [0.195, 0.265] | 0.493 | 12.516 | **<.001** |
|  | **Externalizing ~~**  **Internalizing** | -0.683 (0.047) | [-0.775, -0.591] | -0.758 | -14.600 | **<.001** |
|  | **Externalizing ~~**  **Borderline** | -0.034 (0.016) | [-0.065, -0.003] | -0.151 | -2.093 | **.036** |
|  | Internalizing ~~  Borderline | -0.027 (0.024) | [-0.074, 0.020] | -0.052 | -1.083 | .279 |

**Supplementary Table S5. Parameter estimates of the ST/LT SEM.** Regressions of adjustment variables, intercepts, thresholds and variances are not included for readability (altogether they would add 89 additional parameters), but the full model is available in the supplementary .RData file. Statistically significant terms are highlighted in bold.

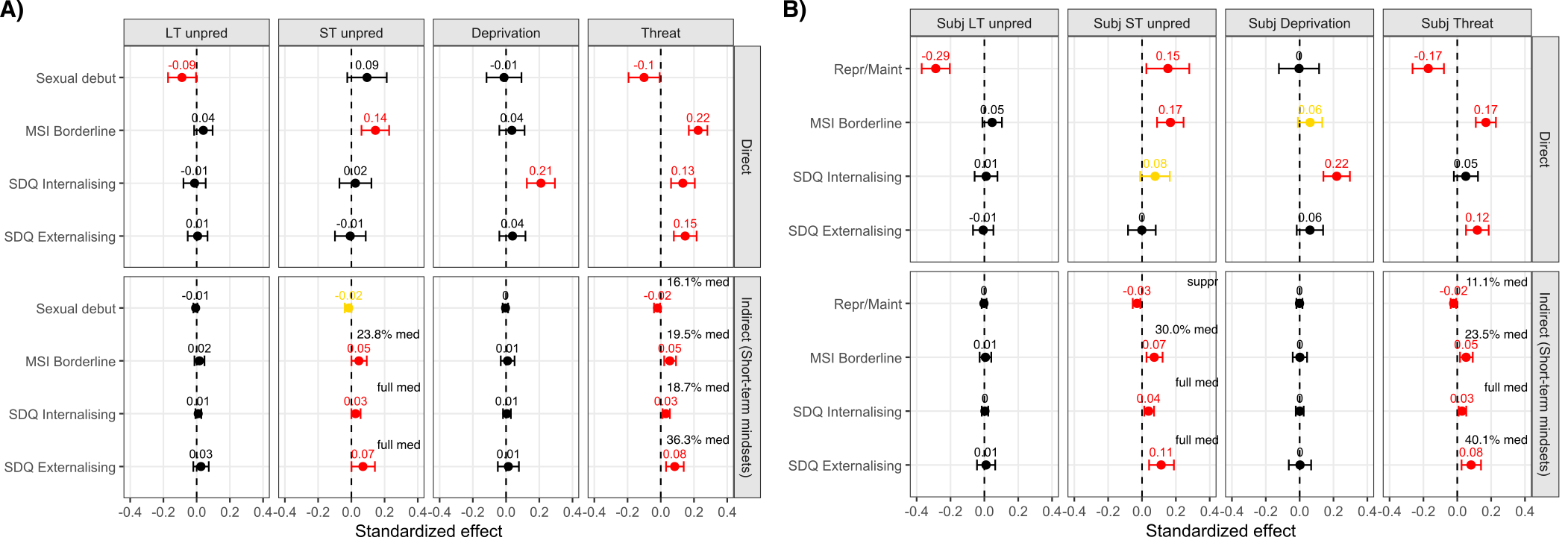
**Supplementary Figure S6. Direct and indirect effects in the sensitivity analysis SEMs.** **A)** Sensitivity analysis using sexual debut only. **B)** Sensitivity analysis using subjective severity weighted adversity. In both panels, estimates of standardized direct and indirect effects of early life adversity dimensions on reproductive/maintenance and mental health. Error bars are Monte Carlo 95% confidence intervals. The proportion of total effect mediated is also shown in case of partial mediation (‘suppr’ and ‘full med’ indicate suppression and full mediation). Parameters significant at α level .05 are in red, Parameters significant at α level .1 are in yellow, parameters not significant are in black.

**Supplementary Text S1**

**No prompt**

I'm now going to give you a list of examples of childhood life experiences. I ask you to create a table where, in one column, you will classify each example of the list as introducing either “stochasticity / short timescale“ or “volatility / long timescale“, and in another column, you will write the corresponding code “1“ or “2“. Limit your answer to one of these two labels:

**Piray and Daw prompt**

I'm going to present a theoretical stance on how the environmental factors to which contemporary humans are exposed can fluctuate:

“It is critical for organisms to infer the true cause of their noisy observations. Take, for example, hiking in the mountains and encountering an unexpected change in weather. It is important to distinguish between regular weather fluctuations, like a passing cloud, and significant weather events, such as an approaching thunderstorm. The appropriate response will depend on accurately assessing the cause of the weather change. The main challenge is that although in both scenarios, the weather is currently less predictable, they should have opposite effects on the hiker’s behavior. If the unexpected weather change is caused by regular fluctuations (referred to as “stochasticity”), the hiker needs to continue with their current course of action. However, if the weather change is caused by a thunderstorm (referred to as “volatility”), the hiker must quickly respond because earlier estimates of the situation quickly become irrelevant.

Significant progress has been made in the field of computational neuroscience regarding our understanding of how organisms learn and adapt in noisy environments. A notable achievement has been the development of models that link error-driven learning to principles of sound statistical inference. This approach of recasting learning as statistical inference has inspired and grounded an influential research program studying neuropsychological systems supporting reinforcement learning and choice.

Classically, error-driven learning relies on two key factors: prediction error, which is the difference between actual and expected outcomes, and the learning rate, which determines the weight assigned to new information. Importantly, the statistical re-interpretation of these rules provides a formal justification for the learning rate, revealing that rather than being an arbitrary free parameter, it should be influenced by the statistical properties of noise in the environment: specifically, both volatility and stochasticity. Higher volatility, indicating rapid environmental changes, reduces the usefulness of old information and requires a higher learning rate. Conversely, higher stochasticity should decrease the learning rate, because more stochastic outcomes provide less information about future outcomes. This perspective has fostered influential research programs focused on constructing hierarchical Bayesian models for learning, which describe how organisms could learn from observations while also inferring their volatility and/or stochasticity. However, despite previous experimental studies demonstrating human adaptability to either volatility or stochasticity (typically manipulated and modeled in isolation from one another, or at least with the second variable changing only incidentally), a more challenging computational question remains unanswered: whether and how organisms infer the true cause of noise, some mixture of volatility or stochasticity, when both factors are unknown and potentially changing. This is challenging because although volatility and stochasticity require opposite adjustments to the learning rate, they are easy to confuse, because they each make observations more noisy, albeit by subtly different patterns.

While both factors have been individually recognized in the field for approximately two decades, the computational challenges that arise when both factors are simultaneously changing have been largely neglected until recently. Volatility, in particular, is an extensively researched concept, with numerous experiments documenting behavioral and neural markers that demonstrate how learning rate increases when volatility is increased. These studies have also explored the disruption of these effects in relation to psychopathologies. Many such studies have systematically manipulated volatility in blocks, with stochasticity changing only incidentally due to (and confounded by) changes in the mean of binomial outcomes. However, even among the few studies that have examined some forms of both volatility and stochasticity systematically played off against one another, they have typically made their dissociation comparatively easy by manipulating volatility by binary jumps of the hidden cause qualitatively different from more graded realization of stochasticity, or have removed the need for computing them by providing explicit information regarding the nature and level of the noise in a specific session. In real-life situations, however, individuals must rely solely on their observations over time to dissociate these factors.

In a recent article, we investigated this issue at the computational level. We showed that although it is necessary for adaptive learning to dissociate whether the cause of noise is volatility or stochasticity, it is computationally difficult to do so because both volatility and stochasticity increase the overall noisiness (i.e., variance) of observations. Nevertheless, it is indeed possible to distinguish between volatility and stochasticity due to a subtle yet crucial difference in their effects on the statistical properties of generated observations. In particular, while both increase the variance of observations, volatility increases their autocorrelation while stochasticity decreases it.

We have shown that taking account of this distinction explains a wide range of neuroscience phenomena and reconciles long-standing competing theories in the associative learning literature on whether surprising outcomes decrease or increase the learning rate by showing that these theories rely on paradigms that, in effect, manipulate either stochasticity or volatility. Furthermore, the distinction between stochasticity and volatility has significant implications for neuropsychiatry. This is because inferring their levels requires disentangling their effects on the observed noise, so that abnormalities in inferring one will tend also to affect the other. Importantly, this implies that much previous work that appeared to demonstrate associations between various psychiatric dysfunction and impaired sensitivity to volatility may instead point to primary or secondary effects mediated by stochasticity (which was typically not manipulated or modeled). We argued that such compensatory tradeoffs may manifest in various disorders that have been previously investigated in the context of volatility, such as learning abnormalities observed in anxiety disorders and following amygdala damage.”

I'm now going to give you a list of examples of childhood life experiences. I ask you to create a table where, in one column, you will classify each example of the list as introducing either “stochasticity / short timescae“ or “volatility / long timescale“, and in another column, you will write the corresponding code “1“ or “2“. Limit your answer to one of these two labels:

**Full prompt**

I'm going to present a theoretical stance on how the environmental factors to which contemporary humans are exposed can fluctuate:

“In this study, we investigate the associations between dimensions of childhood adversity, mental health and LH traits, and test whether they are mediated by short-term mindsets.

We also examine the dimension of unpredictability in more detail, given that it is often found to be even more strongly linked to developmental outcomes than harshness, and at the same time, its conceptualization and measurement remain challenging. There is ample empirical evidence supporting developmental cascade introduced above in which early life unpredictability leads individuals shift to a LH strategy characterized by increased reproductive over maintenance effort by adopting an unpredictability schema and a short-term mindset, leading to risk taking, impulsivity, and externalizing behavioural problems.

However, the literature has not yet reached consensus on how to define and operationalize unpredictability as a construct. Researchers tend to either adopt an ‘ancestral cue’ approach, in which children are assumed to be sensitive to a limited set of specific experiences that served as cues to expected environmental variability in our evolutionary history, or a ‘statistical learning’ approach, which sees children as continuously estimating variability in their environment by more domain-general learning mechanisms. As a result, quite diverse experiences, such as residential changes, subjective retrospective judgments of perceived unpredictability, inconsistent parenting, or income variability have all been used as proxies of unpredictability.

There are some findings suggesting that the timescale and type of uncertainty might be crucial in determining its developmental impact. Here, we tentatively adopt an organizing principle which distinguishes between short-timescale and long-timescale unpredictability. This is partially inspired by the human reinforcement learning (RL) literature, which distinguishes between ‘stochastic’ and ‘volatile’ forms of uncertainty, with important consequences for adaptive behaviour. Stochasticity, or ‘expected uncertainty’, involves random noise around a stable mean (e.g., daily variation in commute time when using public transport), while volatility, or ‘unexpected uncertainty’, signals a true change in outcome contingencies (e.g., a sudden subway closure).

There is extensive empirical evidence that humans can arbitrate between these different forms of uncertainty in their environment, and adopt learning and decision-making styles in an adaptive manner, depending on the degree of inferred volatility and stochasticity. Early life experiences might also be subject to uncertainty classifiable along similar dimensions. For example, day-to-day variability in the time parents come back home is more akin to stochastic unpredictability, whereas an abrupt residential or school change seems closer to volatile unpredictability. Building on this, we also explore whether distinguishing relatively more stochastic, short-timescale from more volatile, long-timescale early life unpredictability experiences has utility for explaining variation in mental health and LH outcomes.“

I'm now going to give you a list of examples of childhood life experiences. I ask you to create a table where, in one column, you will classify each example of the list as introducing either “stochasticity / short timescale“ or “volatility / long timescale“, and in another column, you will write the corresponding code “1“ or “2“. Limit your answer to one of these two labels:
